# Stratification of TAD boundaries identified in reproducible Hi-C contact matrices reveals preferential insulation of super-enhancers by strong boundaries

**DOI:** 10.1101/141481

**Authors:** Yixiao Gong, Charalampos Lazaris, Theodore Sakellaropoulos, Aurelie Lozano, Prabhanjan Kambadur, Panagiotis Ntziachristos, Iannis Aifantis, Aristotelis Tsirigos

## Abstract

The metazoan genome is compartmentalized in megabase-scale areas of highly interacting chromatin known as topologically associating domains (TADs), typically identified by computational analyses of Hi-C sequencing data. TADs are demarcated by boundaries that are largely conserved across cell types and even across species, although, increasing evidence suggests that the seemingly invariant TAD boundaries may exhibit plasticity and their insulating strength can vary. However, a genome-wide characterization of TAD boundary strength in mammals is still lacking. A systematic classification and characterization of TAD boundaries may generate new insights into their function. In this study, we first use fused two-dimensional lasso as a machine learning method to improve Hi-C contact matrix reproducibility, and, subsequently, we categorize TAD boundaries based on their insulation score. We demonstrate that higher TAD boundary insulation scores are associated with elevated CTCF levels and that they may differ across cell types. Intriguingly, we observe that super-enhancer elements are preferentially insulated by strong boundaries, i.e. boundaries of higher insulation score. Furthermore, we perform a pan-cancer analysis to demonstrate that strong TAD boundaries and super-enhancer elements are frequently co-duplicated in cancer patients. Taken together, our findings suggest that super-enhancers insulated by strong TAD boundaries may be exploited, as a functional unit, by cancer cells to promote oncogenesis.

## INTRODUCTION

The advent of proximity-based ligation assays has allowed scientists to probe the three-dimensional chromatin organization at an unprecedented resolution [1, 2]. Hi-C, a high-throughput chromosome conformation variant, has enabled genome-wide identification of chromatin-chromatin interactions [3]. Hi-C has revealed that the metazoan genome is organized in areas of active and inactive chromatin known as A and B compartments respectively [3]. These are further compartmentalized into super-TADs [4], topologically associating domains (TADs) [5–7] and sub-TADs [8], as well as gene neighbourhoods [9].

Several algorithms have been developed to reveal this hierarchical chromatin organization, including Directionality Index (DI) [5], Armatus [10], TADtree [11], Insulation Index (Crane) [12], IC-Finder [13] and others. However, none of these studies has systematically explored the properties of TAD boundaries. Although TADs are seemingly invariant across cell types, mounting evidence suggests that TAD boundaries can vary in strength, ranging from permissive (“weak”) TAD boundaries that allow more inter-TAD interactions to more rigid (“strong”) boundaries that clearly demarcate adjacent TADs [14]. Recent studies have shown that in *Drosophila,* exposure to heat-shock caused local changes in certain TAD boundaries resulting in TAD merging [15]. Another recent study showed that during motor neuron (MN) differentiation in mammals, TAD and sub-TAD boundaries in the *Hox* cluster are not rigid and their plasticity is linked to changes in gene expression during differentiation [16]. It has also been demonstrated that boundary strength is positively associated with the occupancy of structural proteins including CCCTC-binding factor (CTCF) [5]. Despite these advances, no study has yet addressed the issue of boundary strength in mammals and how it may be related to potential boundary disruptions and aberrant gene activation in cancer. Here, we first introduce a new method based on fused two-dimensional lasso [17] in order to improve Hi-C matrix reproducibility. Then, we use the improved Hi-C matrices to: (a) categorize TAD boundaries based on their insulating strength, (b) characterize TAD boundaries in terms of CTCF binding and other functional elements, and (c) investigate potential genetic alterations of TAD boundaries in cancer. We anticipate that our study will help generate new insights into the significance of TAD boundaries.

## MATERIALS AND METHODS

### Initial processing of published high-resolution Hi-C datasets

In order to develop and benchmark a method that improves reproducibility of Hi-C contact matrices, we carefully selected our Hi-C datasets to represent technical variation due to the execution of the experiments by different laboratories and/or the usage of different restriction enzymes. We identified publicly available human Hi-C datasets that fulfilled the following criteria: (i) availability of two biological replicates and (ii) sufficient sequencing depth to robustly identify topologically associating domains (TADs) as described in our TAD calling benchmark study [18]. Specifically, we ensured that our datasets included samples with at least ~40 million intra-chromosomal read pairs and that the Hi-C experiment was performed in biological replicates, either by using one restriction enzyme (HindIII or MboI) (H1 cells and their derivatives [19], K562, KBM7 and NHEK cells [20] and in-house generated CUTLL1), or two enzymes (HindIII or MboI) (GM12878 [20], IMR90 [5, 21]), in order to examine the consistency of predicted Hi-C interactions across different enzymes. Detailed information about the Hi-C datasets, including cell type and GEO accession number, is listed in **Supplementary Table 1**. All datasets were then comprehensively re-analysed using our HiC-bench platform [18]. Briefly, paired-end reads were mapped to the reference genome (hg19) using Bowtie2 [22]. Reads with low mapping quality (MAPQ<30) were discarded. Local alignment of input read pairs was performed, as they often consist of chimeric reads between two (non-consecutive) interacting fragments. Mapped read pairs were subsequently filtered for known artifacts of the Hi-C protocol such as self-ligation, mapping too far from the enzyme’s known cutting sites etc, using GenomicTools [23] *gtools-hic filter* command. More specifically, reads mapping in multiple locations on the reference genome *(multihit),* double-sided reads that mapped to the same enzyme fragment *(ds-same-fragment),* reads whose 5’-end mapped too far *(ds-too-far)* from the enzyme cutting site, reads with only one mappable end *(single-sided)* and unmapped reads *(unmapped),* were discarded. Read pairs that corresponded to regions that were very close (less than 25 kilobases, *ds-too-close)* in linear distance on the genome as well as duplicate read pairs *(ds-duplicate-intra* and *ds-duplicate-inter)* were also discarded. Quality assessment analysis revealed that the samples varied considerably in terms of total numbers of reads, ranging from ~150 million reads to more than 1.3 billion. Mappable reads were over 96% in all samples. The percentages of total accepted reads corresponding to *cis* (ds-accepted-intra, dark green) and *trans* (ds-accepted-inter, light green) also varied widely, ranging from ~17% to ~56%. Duplicate read pairs (*ds-duplicate-intra* and *ds-duplicate-inter,* red and pink respectively), non-uniquely mappable *(multihit,* light blue), single-end mappable *(single-sided;* dark blue) and unmapped reads *(unmapped;* dark purple) were discarded. Self-ligation products *(ds-same-fragment;* orange) and reads mapping too far *(ds-too-far,* light purple) from restriction sites or too close to one another *(ds-too-close;* orange) were also discarded. Only double-sided uniquely mappable *cis (ds-accepted-intra;* dark green) and *trans* (*ds-accepted-inter*; light green) read pairs were used for downstream analysis. Despite the differences in sequencing depth and in the percentages of useful reads across samples, all samples had enough useful reads for TAD detection. Due to the wide differences in sequencing depth, and to ensure fair comparisons of Hi-C matrices in this study, all datasets were down-sampled such that the number of usable intra-chromosomal reads pairs was ~40 million for each replicate. To study the effect of sequencing depth, we also resampled at ~80 million usable intra-chromosomal read pairs. Finally, Hi-C contact matrices were generated using fixed bin sizes at multiple resolutions (5, 20, 40, 60, 80 and 100kb).

### Scaled Hi-C contact matrices, effective length, GC content and mappability

Hi-C contact matrices were scaled by: (a) the total number of (usable) intra-chromosomal read pairs, and (b) the “effective length” of the corresponding pair of interacting bins [24]. More specifically, the scaled Hi-C count corresponding to interactions between the Hi-C matrix bins *i,j* (*y*_*ij*_) is defined by the formula:

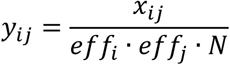

where *X*_*ij*_ is the original number of interactions between the bins *i* and *j, eff*_*i*_ the effective length for the bin *i, eff*_*j*_ the effective length for the bin j, and *N* is the total number of read pairs. For each bin, at each resolution, effective length, GC content and mappability were calculated as originally defined in [25]. In that study, it was demonstrated that the main source of enzyme-specific biases is the effective length. Consequently, we expected that correcting for effective length alone would simultaneously correct for GC content and mappability biases. To verify this, we generated heatmaps showing the association of Hi-C interactions with effective length, GC content and mappability, as described in [26].

### Distance-normalized Hi-C contact matrices

Genomic loci that are further apart in terms of linear distance on DNA tend to give fewer interactions in Hi-C maps than loci that are closer. For intra-chromosomal interactions, this effect of genomic distance should be taken into account. Consequently, the interactions were distance-normalized using an adjusted z-score that was calculated taking into account the mean Hi-C counts for all interactions at a given distance *d* and the corresponding standard deviation. Thus, the z-score for the interaction between the Hi-C contact matrix bins *i* and *j* (*z*_*ij*_) is given the following equation:

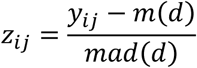

where *y*_*ij*_ corresponds to the number of interactions between the bins *i* and *j, m(d)* to the median number of interactions for distance *d*=|j-i| and *mad(d)* is the robust estimator of the standard deviation of the mean.

### Fused two-dimensional lasso

We used two-dimensional lasso, an optimization machine learning technique widely used to analyse noisy datasets, especially images [17]. This technique is very-well suited for identifying topological domains based on contact maps generated by Hi-C sequencing experiments because topological domains are continuous DNA segments of highly interacting loci that would represent solid squares along the diagonal of Hi-C contact matrices. Topological domains map to squares of different length along the diagonal of the Hi-C contact matrix, but they are not solid as they contain several gaps, i.e. scattered regions on those squares that show little or no interaction. Two-dimensional fused lasso [27] addresses the issue by penalizing differences between neighbouring elements in the contact matrix. This is achieved by the penalty parameter *λ* (lambda), as described in the equation:

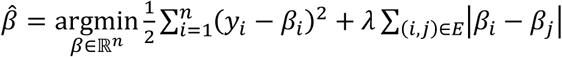

where *y* is the original (i.e. observed) contact matrix, and 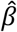 is the optimized contact matrix such that the objective function described above in minimized. *E* describes the neighboring elements of the matrix, i.e. *E* = { (*i, j*), where *i* and *j* are adjacent elements in matrix *β*}.

### Fused two-dimensional lasso packages

We used two R packages that implement fused two-dimensional lasso:

- the flsa R package (https://cran.r-project.org/web/packages/flsa/index.html), for coarse resolutions (up to 20kb) [28]
- for fine resolutions, the more recent and more efficient graph-fused lasso python/C++ package (https://github.com/tansey/gfl) [29].

### Calculation of same-enzyme and cross-enzyme Hi-C matrix reproducibility

We calculated two types of correlation for Hi-C matrices, to evaluate the performance of our method: (a) same-enzyme reproducibility between Hi-C replicates prepared with the same restriction enzyme, (b) cross-enzyme reproducibility between Hi-C replicates prepared with two different enzymes (e.g HindIII/MboI). Hi-C matrix reproducibility was assessed using stratum-adjusted correlation coefficient [30] values, calculated on the filtered, ICE-corrected [31], calCB-corrected [26] and scaled Hi-C contact matrices.

### TAD boundary “ratio” insulation score

Given a potential TAD boundary, we denote the “upstream” region to the left of the boundary as *L,* and the “downstream” region to the right as *R.* The between regions *L* and *R* are denoted as *X*. The “ratio” insulation score is defined as follows:

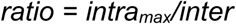

where:

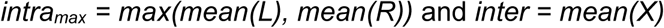

For more details, see [18].

### TAD calling using the “ratio” insulation score

For TAD calling, we first calculated the “ratio” insulation score for each bin at 40kb resolution. Then, TAD boundaries (of size equal to the bin size, i.e. 40kb) were identified as local maxima of the insulation scores across each chromosome. Only insulation scores above a certain cutoff were considered as potential TAD boundaries. The cutoff was determined such that the false discovery rate (FDR) of the identified local maxima was not greater than 10%. The FDR was estimated by applying the same procedure (calculate “ratio” insulation scores and seeking local maxima) on randomized Hi-C matrices. The randomized Hi-C matrices were generated by permuting the original matrix values separately for each “diagonal” of the matrix (i.e. Hi-C interaction values at a given distance between interacting loci), so that the distribution of Hi-C signal as a function of distance between interacting loci was preserved in the randomized matrix. The code is publicly available as part of the HiC-bench distribution.

### Categorization of TAD boundaries based on fused two-dimensional lasso

We applied two-dimensional fused lasso to categorize TAD boundaries based on their strength. The rationale behind this categorization is that topological domains separated by more “permissive” (i.e. weaker) boundaries [32] will tend to fuse into larger domains when lasso is applied, compared to TADs separated by well-defined, stronger boundaries. We indeed applied this strategy and categorized boundaries into multiple groups ranging from the most permissive to the strongest boundaries. The boundaries that were lost when *λ* value was increased from 0 to 0.25, fall in the first category (*λ*=0), the ones lost when *λ* was increased to 0.5, in the second (*λ*=0.25) etc.

### Categorization of TAD boundaries based on insulation scores

We stratified TAD boundaries into five categories (I through V) of equal size according to their insulation score, independently in each Hi-C dataset used in this study. Category I contained TAD boundaries with the lowest insulation scores and category V contained those with the highest. Before calculating insulation scores, we first processed the Hi-C matrices using ICE, calCB and scaling and then applied fused 2D lasso (with optimal *λ*). Then, TAD calling and TAD boundary insulation score calculations were performed using our “ratio” or the “crane” method and the boundaries were classified into five equal-size categories, as described above.

### Analysis of CTCF and H3K27ac ChIP-seq data

All ChIP-seq data was uniformly processed using the HiC-bench platform [18]. Raw sequencing files were aligned using bowtie2 version 2.3.1 with standard parameters. Only uniquely mapped reads were selected for downstream analysis. PCR duplicates were removed using Picad-tools version 1.88. Macs2 version 2.0.10.20131216 were used to call narrow peaks for CTCF and broad peaks for H3K27ac with default parameters.

### Association of CTCF levels with boundary strength categories

We obtained CTCF ChIP-sequencing data for the cell lines utilized in this study (with the exception of KBM7 for which no publicly available dataset was available, see **Supplementary Table 1** for details). Total CTCF levels (i.e. aggregated peak intensities from potentially multiple CTCF peaks) at each TAD boundary were calculated and their normalized distributions for each boundary category (weak to strong) were plotted in boxplots in order to demonstrate the association of increased boundary strength with increased levels of CTCF binding. We performed this analysis separately for TSS-only and non-TSS CTCF binding sites. The rationale behind these separate analyses was based on the observation that several TAD boundaries, especially strong boundaries, contain TSSs. Statistical significance was assessed using paired two-sided Wilcoxon rank sum test. The boxplots represent the distribution of values (normalized CTCF levels) across the Hi-C samples used in this study to define the five categories of TAD boundaries. Detailed information about the CTCF ChIP-seq datasets, including cell type and GEO accession number, is made available in **Supplementary Table 1**.

### Association of boundary strength categories with proximity to super-enhancers

Super-enhancers were called using H3K27ac ChIP-seq data from GEO, ENCODE and in-house generated data. Detailed information about the H3K27ac ChIP-seq datasets, including cell type and GEO accession number, is made available in **Supplementary Table 1**. Reads were first aligned with Bowtie2 v2.3.1 [22] and then HOMER v4.6 [33] was used to call super-enhancers, all with standard parameters. For each super-enhancer in each sample, we identified the corresponding TAD and its TAD boundaries. We then calculated (per sample) the percentage of super-enhancers that are surrounded by boundaries belonging in each boundary category. Statistical significance was assessed using paired two-sided Wilcoxon rank sum test. The boxplots represent the distribution of values (fraction of super-enhancers in proximity to TAD boundary categories I through V) across the Hi-C samples used in this study.

### Pan-cancer analysis of deletions, duplications and co-duplications of enhancer/super-enhancer elements with TAD boundaries

Deletion and co-duplication data were downloaded from ICGC [34]. Then, deletions and co-duplications were categorized based on their size ranging from 250kb to 10Mb. This data was combined with boundary strength data (from the cell lines included in this study) and the closest boundaries to each structural variant were identified using BEDTools [35]. Data for super-enhancers were downloaded from the super-enhancer archive (SEA) [36], whereas enhancer data were downloaded from FANTOM [37]. Then, the fraction of boundaries or enhancer/super-enhancer elements was normalized with the total size of the corresponding structural variation data (deletions or tandem duplications) and plotted against boundary strength. Statistical significance was assessed using paired two-sided Wilcoxon rank sum test. The boxplots represent the distribution of values (fraction of boundaries or enhancer/super-enhancers in proximity to TAD boundary categories I through V) across the Hi-C samples used in this study.

## RESULTS

### Analysis workflow

The overall workflow, including our benchmark strategy and downstream analysis, is summarized in **Figure 1**. Initial alignment and filtering of the collected Hi-C sequencing datasets was performed with HiC-bench [18] (see Methods for details). Quality assessment analysis revealed that the samples varied considerably in terms of total numbers of reads, ranging from ~150 million reads to more than 1.3 billion (**Supplementary Figure 1a**). Mappable reads were over 96% in all samples. The percentages of total accepted reads corresponding to *cis* (ds-accepted-intra, dark green) and *trans* (ds-accepted-inter, light green) (**Supplementary Figure 1b**) also varied widely, ranging from ~17% to ~56%. The characteristic drop of average Hi-C signal as a function of distance between interacting loci was observed (**Supplementary Figure 1c**). The main part of analysis starts with unprocessed Hi-C contact matrices (“filtered” matrices). We then generate processed Hi-C matrices using ICE “correction” [31], our “scaling” approach (see Methods) and calCB [26]. Finally, fused two-dimensional lasso is applied on the processed Hi-C matrices. Matrix reproducibility between biological replicates is assessed across samples for a variety of parameters, for example, resolution, distance between interacting loci, sequencing depth, etc, using stratum-adjusted correlation coefficients [30]. Finally, downstream analysis, involves the characterization of TAD boundaries based on their insulating strength, the enrichment in CTCF binding, proximity to repeat elements and super-enhancers, and, finally, their genetic alterations in cancer.

**Figure 1.**
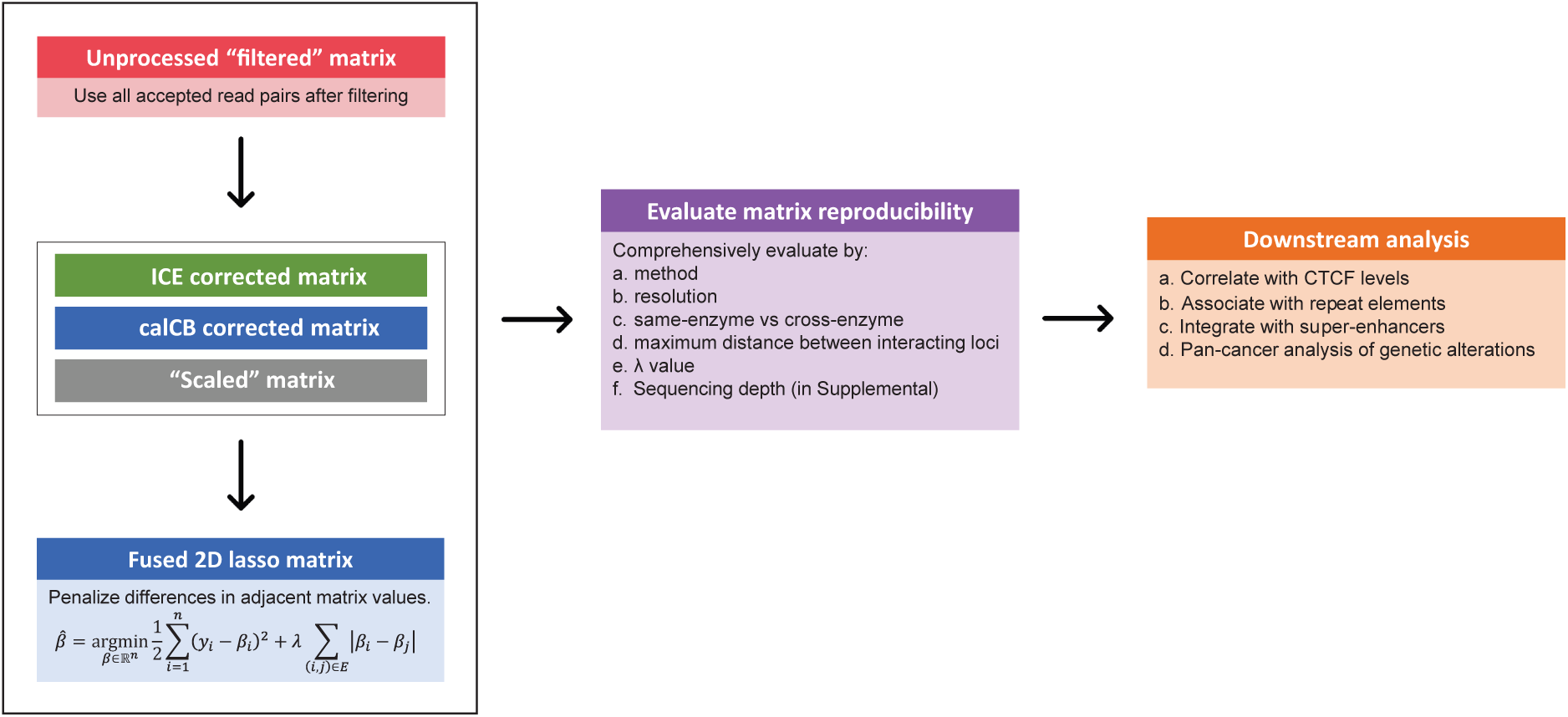
Overall workflow and benchmarking strategy. Our analysis starts with unprocessed Hi-C contact matrices. We then generate processed Hi-C matrices using ICE “correction”, our “scaling” approach and calCB. Fused two-dimensional lasso is applied on the processed Hi-C matrices. Matrix reproducibility between biological replicates is assessed across samples for a variety of parameters using stratum-adjusted correlation coefficients [30]. Finally, downstream analysis, involves the characterization of TAD boundaries based on their insulating strength, the enrichment in CTCF binding, proximity to repeat elements and super-enhancers, and, their genetic alterations in cancer.

### Assessment of same-enzyme and cross-enzyme reproducibility of Hi-C contact matrices

Hi-C is prone to biases and multiple algorithms have been developed for Hi-C bias correction, including probabilistic modelling methods [24], Poisson or negative binomial normalization [25], calCB which corrects for copy number variation [26] and the widely used Iterative Correction and Eigenvalue decomposition method (ICE) [31] which assumes “equal visibility” of genomic loci. A similar iterative method named Sequential Component Normalization was introduced by Cournac *et al.* [38]. Additional efficient correction methods have been developed to handle high-resolution Hi-C datasets [39]. However, estimating highly reproducible Hi-C contact maps remains a challenging task [40], especially at high resolutions, as we also demonstrate below. Specifically, we focused on multiple factors that may play an important role on reproducibility: *first,* we separately considered biological replicates of Hi-C libraries generated with the same or different restriction enzymes; *second,* we studied the impact of Hi-C matrix resolution (i.e. bin size); *third,* we assessed reproducibility as a function of the distance of interacting loci pairs; *fourth,* we studied the impact of sequencing depth. Stratum-adjusted correlation coefficients (SCC) were calculated for each pair of replicates (same- or cross-enzyme) on Hi-C contact matrices estimated by four methods: (i) naïve filtering (i.e. matrix generation by simply using double-sided accepted intra-chromosomal read pairs from **Supplementary Figure 1a**), (ii) iterative correction (ICE) which has been demonstrated to improve cross-enzyme correlation, (iii) calCB which corrects for known Hi-C biases as well as for copy number variation and (iv) our own scaling method which also corrects for effective length, GC content and mappability (see **Supplementary Figure 2a,b** and Methods for details). The results of our benchmark analysis are summarized in **Supplementary Figure 3**: the left panel summarizes the correlations between replicates generated by the same restriction enzyme, whereas the right panel the correlations between replicates generated by a different restriction enzyme. In both scenarios, as expected, reproducibility drops quickly as finer resolutions (from 100kb to 20kb) are considered. The same conclusion applies for increasing distance (from 2.5Mb to 10Mb) between interacting loci, demonstrating that long-range interactions require ultra-deep sequencing (beyond what is currently available in most of the datasets in this study) in order to be detected reliably. To elaborate on this point, we repeated the analysis after resampling at higher sequencing depth (**Supplementary Figure 4**). Both conclusions hold true with the new sequencing depth and are independent of the Hi-C contact matrix estimation method. From this benchmarking study, we conclude that reproducibility of Hi-C contact matrices across biological replicates is not ideal and that there is a need for computational methods to improve it. In the next sections, we focus on improving the reproducibility of the Hi-C contact matrices within the context of TADs, as most of the DNA-DNA interactions occur within these domains. Since TAD sizes typically range from 200Kb-2.5Mb (>92% of all TADs identified in our Hi-C datasets), and, as demonstrated in **Supplementary Figure 3** and **Supplementary Figure 4**, stratum-adjusted correlation coefficients between biological replicates of Hi-C contact matrices drop dramatically beyond 2.5Mb, we restrict our subsequent analyses to distances up to 2.5Mb.

### Fused lasso improves same-enzyme and cross-enzyme correlations of Hi-C contact matrices

Motivated by the poor performance of all methods at fine resolutions and by the observation of a trade-off between cross-enzyme and same-enzyme reproducibility when correcting for enzyme-related biases, we decided to utilize a machine learning denoising method, fused two-dimensional lasso [27], to improve the reproducibility of Hi-C contact matrices. Briefly, two-dimensional fused lasso introduces a parameter *λ* which penalizes differences between neighboring values in the Hi-C contact matrix (see Methods for details). The effect of parameter *λ* is demonstrated in **Figure 2a** where we show an example of the application of fused two-dimensional lasso on a Hi-C contact matrix focused on an 8Mb locus on chromosome 8 (chr8:124700000-132700000) for different values of parameter *λ*. To evaluate the performance of fused lasso, we calculated same-enzyme and cross-enzyme stratum-adjusted correlation coefficient (SCC) values between Hi-C contact matrices generated from different replicates. SCC values were calculated either for iteratively-corrected (ICE), calCB-corrected or scaled Hi-C contact matrices (at different *λ* values) and compared to the naïve filtering approach. The results for same enzyme, are summarized in **Figure 2b**. Increasing *λ* improves reproducibility independent of resolution, restriction enzyme and bias-correction method, demonstrating the robustness of our approach. Similarly, fused two-dimensional lasso improves the reproducibility of contact matrices in the cross-enzyme case, as demonstrated in **Figure 2c**. The same analysis was performed at lower sequencing depth with similar results (**Supplementary Figure 5**). Next, we explored the effect of fused 2D lasso on Hi-C matrices of fine resolutions, and we tested whether the lasso approach can improve detection of specific DNA-DNA interactions. For this analysis, we used 5kb bins to compute the interaction matrix. To compensate for distance-related biases in Hi-C matrices (**Supplementary Figure 1c**), we normalized the interaction strength for every distance/diagonal using a robust version of z-score (see Methods for details). Then, we applied the Graph-Fused Lasso implementation of fused 2D lasso [29], which scales better than the Fused Lasso Signal Approximator (flsa) [28] used for coarse resolutions. Since available Hi-C datasets lack biological replicates of ultra-deep sequenced samples, we evaluated our method by testing whether it could recover the 5kb loops identified in [20] in a single biological sample of GM12878, the most deeply sequenced sample in this study (~3 billion read pairs of which ~900 million intra-chromosomal read pairs passed our filtering criteria). As a recovery metric, we used the fraction of the reported loops within the top interactions as ranked by our fused lasso approach. We observed that by tuning the *λ* parameter we improved this metric by an 8% relative improvement **(Supplementary Figure 6a**). For the optimal *λ*, our method ranked most of the known loops (~90%) in the top 10% of measured interactions (~79% in the top 5% of all measured interactions). We also evaluated the sensitivity of our approach to subsampling. In particular, we re-computed the interaction matrices using 200, 400, 600, 800 million intra-chromosomal read pairs, rerun the analysis and obtained relative improvements of 9%, 14%, 22% and 32% respectively **(Supplementary Figure 6a**). Performance of the graph-fused lasso algorithm as a function of chromosome size is presented in **Supplementary Figure 6b** (peak memory consumption) and **Supplementary Figure 6c** (execution time).

**Figure 2.**
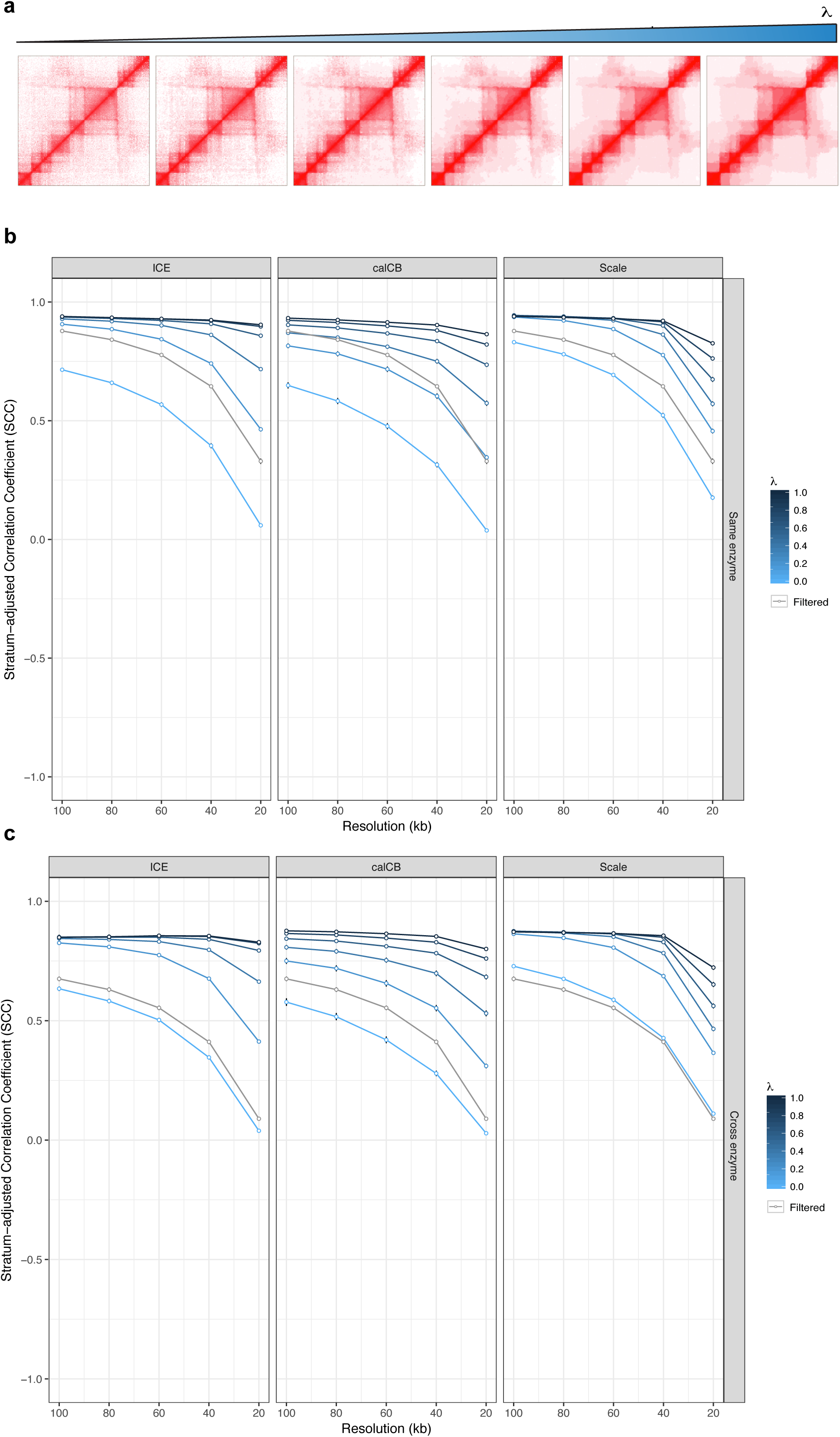
Fused two-dimensional lasso improves reproducibility of Hi-C contact matrices*(high sequencing depth)* **(a)** Example of application of fused two-dimensional lasso on a Hi-C contact matrix focused on a 8Mb locus on chromosome 8 for different values of parameter *λ* (top panel=original matrix; bottom panel=distance-normalized matrix), **(b)** Stratum-adjusted correlation coefficient values are improved by increasing the value of fused lasso parameter *λ* for matrices estimated with ICE, calCB and our simple scaling method (same enzyme), **(c)** Stratum-adjusted correlation coefficient values are improved by increasing the value of fused lasso parameter *λ* for matrices estimated with ICE, calCB and our simple scaling method (cross enzyme). As a baseline control, stratum-adjusted correlation coefficients of Hi-C contact matrices generated by the naive filtering method are marked by the gray line in each panel.

### Fused lasso preserves cell-type specificity of Hi-C contact matrices

Although fused 2D lasso improves reproducibility of Hi-C matrices between biological replicates, there is a possibility that this is achieved at the expense of losing cell-type specificity. To test this, we compared the effect of *λ* on the reproducibility between biological replicates *(intra-cell-type)* to its effect on the stratum-adjusted correlation coefficients between unrelated samples in our Hi-C dataset collection *(inter-cell-type).* For this test, we chose to focus on the collection of H1 stem cell Hi-C replicates and their derivatives generated by the Ren lab [19], so that we could assess the effect of smoothing on subtle cell-type specific differences in experiments performed in a single lab. Hi-C matrices were distance-normalized (similar to [41], see Methods for details) to account for the dependence of the Hi-C signal on the distance between interacting loci. The results of this analysis are presented in **Figure 3**: although both *intra-cell-type* and *inter-cell-type* stratum-adjusted correlation coefficients increase by *λ* (**Figure 3a**), the difference between *intra-cell-type* and *inter-cell-type* correlation coefficients also increases (**Figure 3b**), suggesting that fused 2D lasso actually preserves cell type specificity of Hi-C contact matrices, a behavior that is consistent independent of the matrix “correction” method. Nevertheless, some “correction” methods appear to work better than others in combination with lasso. In addition, we also evaluated an alternative “smoothing” method, two-dimensional mean filter smoothing, recently made available as part of the HiCRep package [30]. In **Figure 3c**, we show the results of the comparison of the three correction methods in combination with the smoothing techniques using two metrics: preservation of cell type specificity (x axis) and intra-cell-type reproducibility (y axis). The main conclusions from this comparison are: (a) smoothing (lasso or mean filter) improves both metrics independent of the correction method, and (b) fused lasso performs slightly better than mean filter smoothing in preserving cell type specificity, while it behaves slightly worse in improving intra-cell-type specificity. In **Figure 3d**, we further demonstrate the trade-off between *intra-cell-type* and *inter-cell-type* metrics when using 2D lasso or 2D mean filtering.

**Figure 3.**
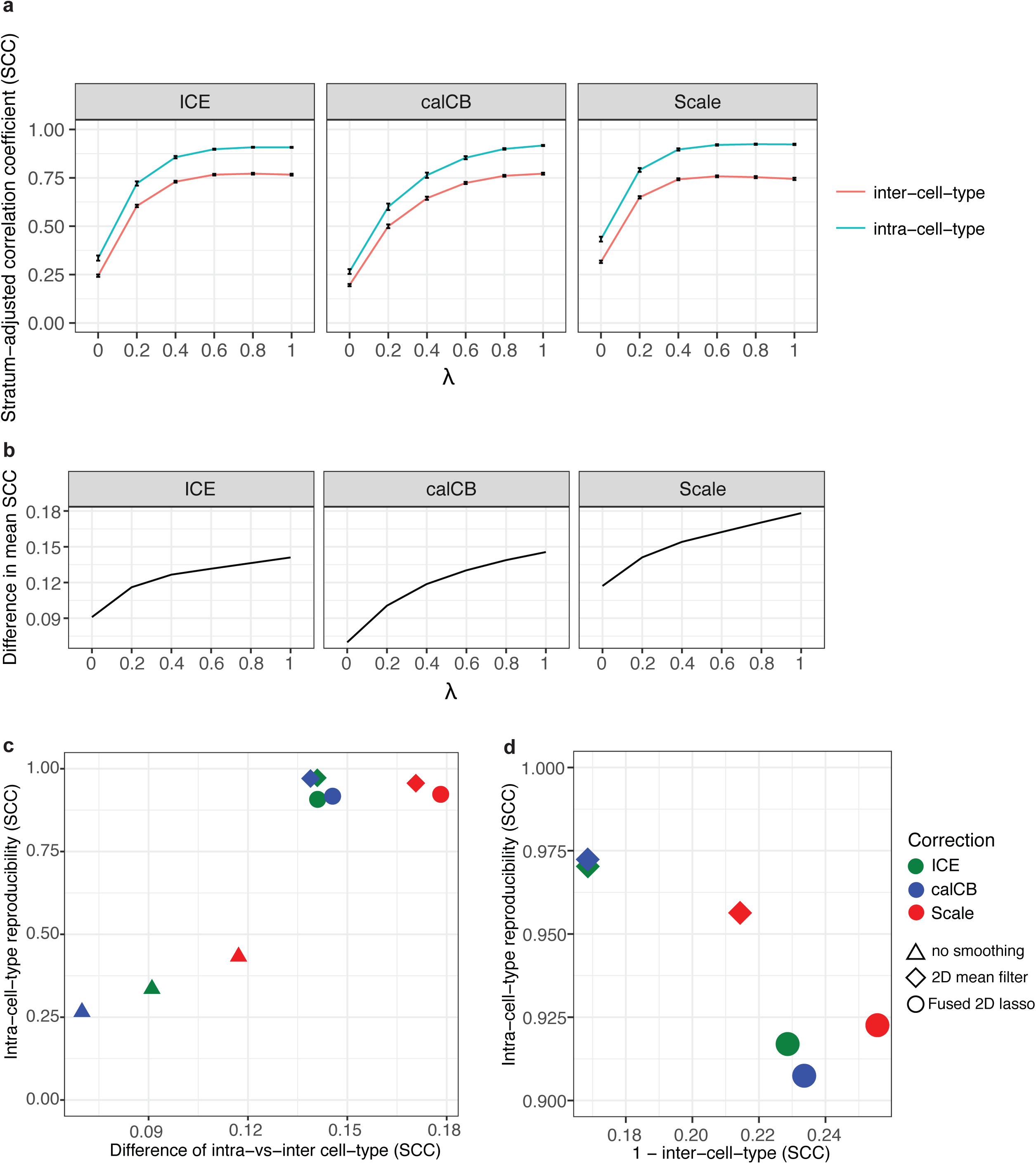
Fused lasso preserves cell-type specificity of Hi-C contact matrices. **(a)** Effect of fused lasso on stratum-adjusted correlation coefficient for the case of *intra-cell-type* (cyan) and *inter-cell-type* (orange) comparisons. Matrices of 40kb resolution were used for the analysis. The Hi-C matrices were processed with ICE, calCB or scaling matrix correction methods, **(b)** Difference in mean stratum-adjusted correlation coefficient between *intra-cell-type* and *inter-cell-type* sample comparisons, **(c)** Comparison of Hi-C matrix “correction” and smoothing methods in terms of preservation of cell type specificity (x axis) and *intra-cell-type* reproducibility (y axis), **(d)** Comparison of Hi-C matrix “correction” and smoothing methods in terms of *(1 - inter-cell-type) (x* axis) and *intra-cell-type (y* axis) stratum-adjusted correlation coefficients.

### Fused lasso reveals a nested TAD hierarchy linked to TAD boundary insulation scores

After demonstrating that parameter *λ* improves reproducibility of Hi-C contact matrices independent of the bias-correction method, we hypothesized that increased values of *λ* may also define distinct classes of TADs with different properties. For this reason, we now allowed *λ* to range from 0 to 5. We then identified TADs at multiple *λ* values using HiC-bench, and we observed that the number of TADs is monotonically decreasing with the value of *λ* (**Supplementary Figure 7a**), suggesting that by increasing *λ*, we are effectively identifying larger TADs encompassing smaller TADs detected at lower *λ* values. Indeed, when comparing TAD boundaries detected at successive *λ* values, we found that higher *λ* values produced TAD boundaries that are almost a strict subset of TAD boundaries produced at lower *λ* values (~94% overlap when considering only the exact bin as a true common TAD boundary, and ~98% when TAD boundaries are allowed to differ by at most one bin between TADs generated for successive *λ* values). Equivalently, certain TAD boundaries “disappear” as *λ* is increased. Therefore, we hypothesized that TAD boundaries that disappear at lower values of *λ* are weaker (i.e. lower insulation score), whereas boundaries that disappear at higher values of *λ* are stronger (i.e. higher insulation score). To test this hypothesis, we identified the TAD boundaries that are “lost” at each value of *λ*, and generated the distributions of the insulation scores for each *λ* across samples. As insulation score, we used the Hi-C “ratio” score (see Methods), which was shown to outperform other TAD calling methods [18]. Indeed, as hypothesized, TAD boundaries lost at higher values of parameter *λ* are associated with higher TAD insulation scores (**Supplementary Figure 7b**).

### Stratification of TAD boundaries by insulating score reveals an association with enriched CTCF levels

Motivated by the observation that with increasing *λ*, weaker TAD boundaries are not detected, we decided to explore in depth the properties of TAD boundaries with respect to their insulation score. To this end, we stratified TAD boundaries into five categories (I through V) of equal size according to their insulation score, independently in each Hi-C dataset used in this study. As shown in **Figure 4a**, we first processed the Hi-C matrices using ICE, calCB and scaling and applied fused 2D lasso with “optimal” *λ*, defined as the *λ* value beyond which no statistically significant improvement on the reproducibility is observed. The statistical significance was assessed using a Wilcoxon test between the distributions of stratum-adjusted correlation coefficients across chromosomes in given sample for successive *λ* values. The procedure is demonstrated using an IMR90 replicate as an example (**Supplementary Figure 7c**). Then, TAD calling and TAD boundary insulation score calculations were performed using our “ratio” method (see Methods for details) and the boundaries were classified into five equal-size categories, as mentioned above. A heatmap representation including all TAD boundaries and their associated boundary strength category across all samples is depicted in **Figure 4b** (*“NA”* corresponds to lack of boundary, as it is possible that boundaries called in certain samples are not present in others). Unbiased hierarchical clustering correctly grouped replicates and related cell types independent of enzyme biases or batch effects related to the lab that generated the Hi-C libraries, suggesting that TAD boundary strength can be used to distinguish cell types. Equivalently, this finding suggests that, although TAD boundaries have been shown to be largely invariant across cell types, a certain subset of TAD boundaries may exhibit varying degrees of strength in different cell types. Also, as expected, TAD boundary strength was found to be positively associated with CTCF levels, suggesting that stronger CTCF binding confers stronger insulation. Since we noticed that several TAD boundaries contain TSSs, this analysis was done separately for TSS-only CTCF peaks (**Figure 4c**) and for all CTCF peaks (see below). Both approaches revealed the same trend, with the exception of the class of strongest boundaries (category V), where CTCF levels in TSS regions were significantly higher compared to non-TSS regions, suggesting that the strongest boundaries are formed by CTCF-mediated loops at gene promoters. Alternatively to our “ratio” insulation score, we repeated our analysis using the insulation score generated by the “crane” TAD calling algorithm [12]. A comparative analysis with between “ratio” and “crane” is shown in **Figure 4d**, where it appears that ratio-generated insulation scores better associate with CTCF levels. In the interest of robustness, we performed the same analyses for all preprocessing methods, at both low and high sequencing depth, for both “ratio” and “crane” insulation scores **(Supplementary Figure 8 and Supplementary Figure 9** respectively), for TSS-only CTCF peaks **(Supplementary Figure 8a** and **Supplementary Figure 9a**), as well as for all CTCF peaks **(Supplementary Figure 8b** and **Supplementary Figure 9b**). Finally, SINE elements have also been shown to be enriched at TAD boundaries [5], and besides confirming this finding, we now demonstrate that Alu elements (the most abundant type of SINE elements) are enriched at stronger TAD boundaries (**Supplementary Figure 10**, top-left panel). A comprehensive analysis of all major repetitive element subtypes (downloaded from the UCSC Genome Browser [42]) can be found in **Supplementary Figure 10**.

**Figure 4.**
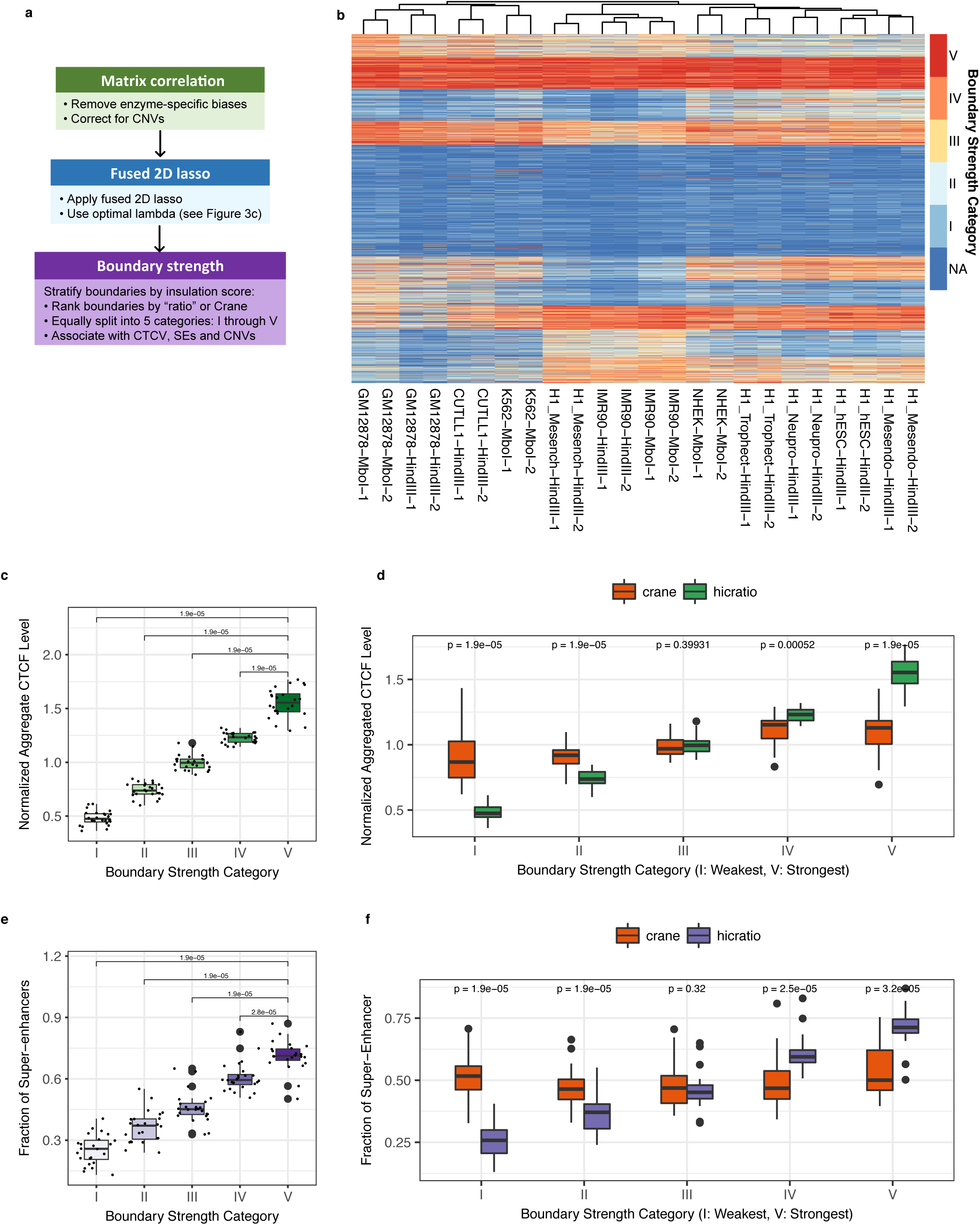
Classification and characterization of TAD boundaries according to insulation score. **(a)** Workflow of stratification of TAD boundaries by insulating score, **(b)** Heatmap representation of TAD boundary insulation strength across samples; hierarchical clustering correctly groups replicates and related cell types independent of enzyme biases or batch effects related to the lab that generated the Hi-C libraries (detailed information about all Hi-C datasets and their cell types is included in **Supplementary Table 1**), **(c)** TAD boundary strength is associated with CTCF levels, **(d)** Comparison of the association of “ratio” vs “crane” insulation scores with CTCF levels, **(e)** Fraction of super-enhancer elements in the vicinity of boundaries of variable strength, **(f)** Comparison of the association of “ratio” vs “crane” insulation scores with respect to proximity to super-enhancers. All statistical tests are paired two-sided Wilcoxon rank sum tests between distributions defined across samples (each sample is a dot in the boxplots).

### Super-enhancers are preferentially localized within TADs demarcated by at least one strong boundary

We then explored what type of functional elements are localized within TADs demarcated by strong TAD boundaries. Specifically, we tested super-enhancers identified in matched samples (see Methods for details). Super-enhancers are key regulatory elements thought to be defining cell identity [9, 43], and are usually found near the center of TADs [44]. Our analysis determined that they are significantly more frequently localized within TADs insulated by at least one strong TAD boundary (**Figure 4e**). Further analysis revealed that, super-enhancers are 2.94 times more likely to be insulated by strong boundaries (categories IV or V) in both the upstream and downstream directions, compared to being insulated by weak boundaries (categories I or II) in both directions. A comparison with TAD boundary classification using “crane” insulation scores demonstrated that “ratio” insulation scores are more significantly associated with proximity to super-enhancers (**Figure 4f**). A similar robustness analysis as the one presented above for CTCF was also performed for super-enhancers **(Supplementary Figure 8c** and **Supplementary Figure 9c** for “ratio” and “crane” insulation scores respectively). Taken together, our findings suggest that, because of their significance in gene regulation, super-enhancers should only target genes confined in the “correct” TAD or neighborhood, while remaining strongly insulated from genes in adjacent TADs. This is conceivably achieved by the strong TAD boundaries we have identified in this study.

### Strong TAD boundaries are co-duplicated with super-enhancers in cancer patients

To further investigate the importance of variable boundary strength, we asked whether TAD boundaries are prone to genetic alterations in cancer. To this end, we mined structural variants released by the International Cancer Genome Consortium (ICGC) [45]. A summary of the reported variant types across all cancer types available on ICGC, is presented in **Supplementary Figure 11**. First, for each focal (up to 1Mb) deletion event, we identified the TAD boundaries closest to the breakpoints, and calculated the frequency of deletions by boundary strength. We observed that the frequency of deletions monotonically decreased with increasing boundary strength (**Figure 5a**). This suggests that strong TAD boundaries are less frequently lost in cancer, as they may “safeguard” functional elements that are necessary for proliferation. By contrast, the frequency of tandem duplications (up to 1Mb) increased with increasing boundary strength (**Figure 5b**). Both results were robust to various cutoffs on the sizes of the structural variants, within the usual range of TAD sizes (from 250kb to 2.5Mb). Then, to further clarify the connection between super-enhancers, strong TAD boundaries and cancer, we studied tandem duplication events where super-enhancers (obtained from a publicly available collection of super-enhancers [36]) are co-duplicated with adjacent strong boundaries. As demonstrated in **Figure 5c**, super-enhancers are indeed co-duplicated with strong TAD boundaries. Co-duplication of strong boundaries and super-enhancers was statistically significantly more frequent than that of strong boundaries and regular enhancers. This suggests that, in cancer, not only are strong boundaries protected from deletions, but they are also co-duplicated with super-enhancer elements. A robustness analysis similar to the one performed for CTCF and super-enhancers is presented in **Supplementary Figure 12** demonstrating that our findings are consistent for low and high sequencing depth. Finally, we present an example of a co-duplication of a super-enhancer with a strong boundary in **Figure 5d**: *MYC,* a well-known oncogene that is typically overexpressed in cancer, is localized next to a strong TAD boundary and is co-duplicated with the boundary as well as with several proximal super-enhancers.

**Figure 5.**
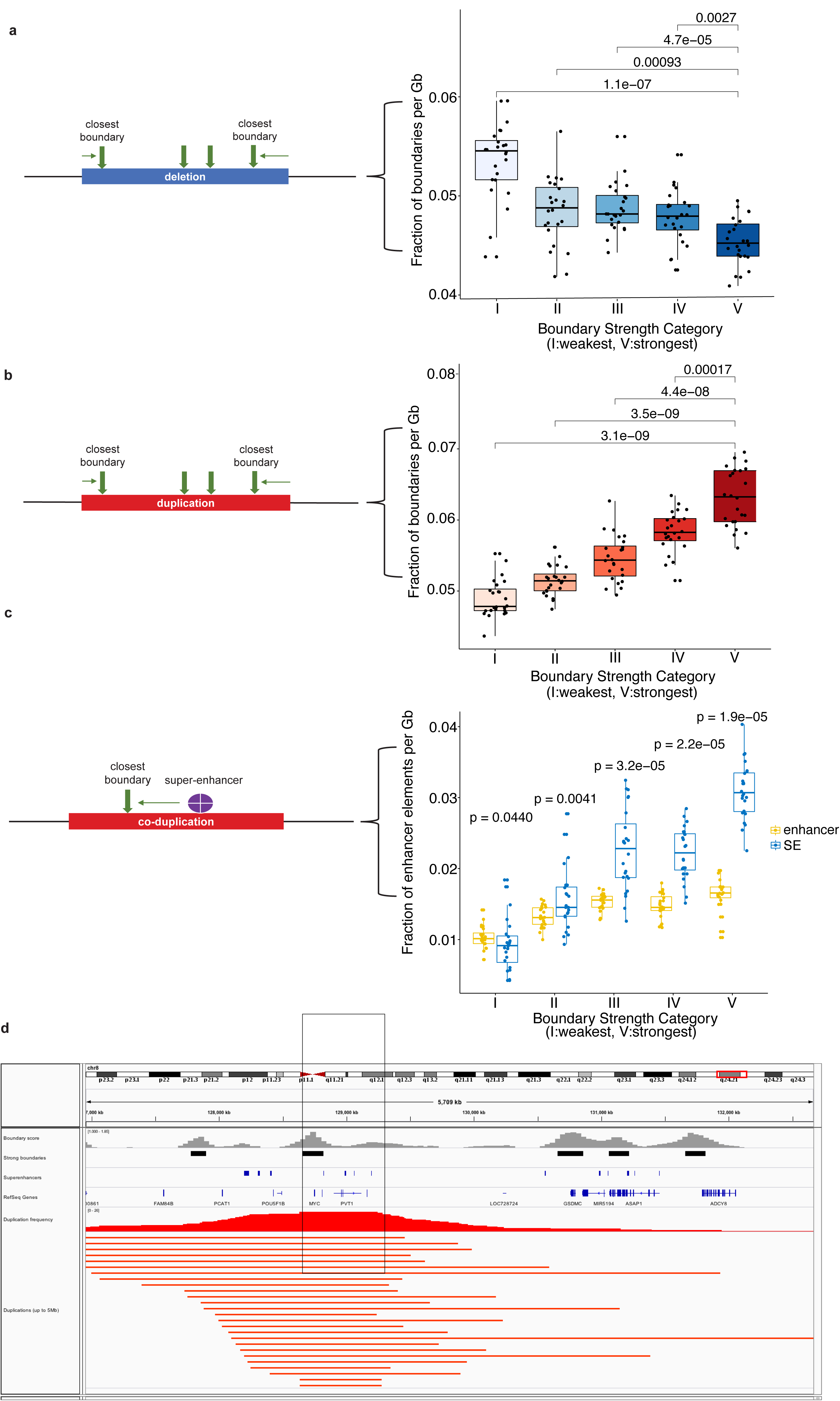
Pan-cancer analysis of strong vs weak TAD boundaries. **(a)** Schematic of pan-cancer analysis (left panel) and classification of focally deleted boundaries in cancer according to their strength (right panel), **(b)** Schematic of pan-cancer analysis (left panel) and classification of focally duplicated boundaries in cancer according to their strength (right panel), **(c)** Schematic of pan-cancer analysis (left panel) and co-duplications of TAD boundaries with regular enhancers and super-enhancers in cancer (right panel), **(d)** Snapshot of the *MYC* locus: a strong boundary (black bar) is frequently co-duplicated with *MYC* and potential super-enhancers in cancer patients (highlighted area). IGV tracks from top to bottom: average insulation score across cell types (gray), strong boundaries (black bars), super-enhancer track from SEA (blue bars), RefSeq genes, duplication frequency (red graph) and ICGC patient tandem duplications (red bars). All statistical tests are paired two-sided Wilcoxon rank sum tests between distributions defined across samples (each sample is a dot in the boxplots).

## DISCUSSION

Multiple recent studies have revealed that the metazoan genome is compartmentalized in boundary-demarcated functional units known as topologically associating domains (TADs). TADs are highly conserved across species and cell types. A few studies, however, provide compelling evidence that specific TADs, despite the fact that they are largely invariant, exhibit some plasticity. Given that TAD boundary disruption has been recently linked to aberrant gene activation and multiple disorders including developmental defects and cancer, categorization of boundaries based on their strength and identification of their unique features becomes of particular importance. In this study, we first developed a method based on fused two-dimensional lasso in order to improve Hi-C contact matrix reproducibility between biological replicates. Then, we categorized TAD boundaries based on their insulating score. Our analysis demonstrated that: (a) using fused 2D lasso, we can improve the reproducibility of Hi-C contact matrices irrespective of the Hi-C bias correction method used, and (b) using our “ratio” insulation score, we can successfully identify boundaries of variable strength and that strong boundaries exhibit certain expected features, such as elevated CTCF levels. By performing an integrative analysis of boundary strength with super-enhancers in matched samples, we observed that super-enhancers are preferentially insulated by strong boundaries, suggesting that super-enhancers and strong boundaries may represent a biologically relevant entity. Motivated by this observation, we examined the frequency of structural alterations involving strong boundaries and super-enhancers. We found that not only strong boundaries are “protected” from deletions, but, more importantly, they are co-duplicated together with super-enhancers. Recently, it has been shown that genetic or epigenetic alterations near enhancers may lead to aberrant activation of oncogenes [46–49]. Our results, expand on these studies by highlighting a previously unknown connection between strong TAD boundaries, super-enhancers and tandem duplication events in cancer.

## AUTHOR CONTRIBUTIONS

AT conceived this study. AT, AL and PK proposed the use of lasso optimization on Hi-C contact matrices. AT designed the computational experiments. YG extended HiC-bench (HiCRep, mean 2D filter smoothing and calCB). YG, CL and AT performed computational analyses and generated figures. TS performed analysis of Hi-C matrices at fine resolutions using graph-fused lasso. PN performed the CUTLL1 Hi-C experiments. PN and IA offered biological insights and helped with the interpretation of Hi-C data. AT wrote the Methods and Results with help from CL. CL wrote the Introduction and Discussion with help from AT. All authors read and approved the final manuscript.

## ACKNOWLEDGMENTS

We would like to thank all members of the Tsirigos and Aifantis Laboratories for critical evaluation of the manuscript, and Andreas Kloetgen who provided guidance on identifying specific interactions at fine resolutions. We would like to thank the Applied Bioinformatics Laboratories (ABL) at the NYU School of Medicine for providing bioinformatics support and helping with the analysis and interpretation of the data. This work has used computing resources at the NYU High Performance Computing Facility (HPCF). We also thank the Genome Technology Center (GTC) for expert library preparation and sequencing. This shared resource is partially supported by the Cancer Center Support Grant, P30CA016087, at the Laura and Isaac Perlmutter Cancer Center.

## FUNDING

The study was supported by the American Cancer Society [RSG-15-189-01-RMC to AT] and a Leukemia & Lymphoma Society New Idea Award [8007-17 to AT]. NYU Genome Technology Center (GTC) is a shared resource, partially supported by the Cancer Center Support Grant [P30CA016087] at the Laura and Isaac Perlmutter Cancer Center.

## TABLE AND FIGURE LEGENDS

**Supplementary Figure 1.**
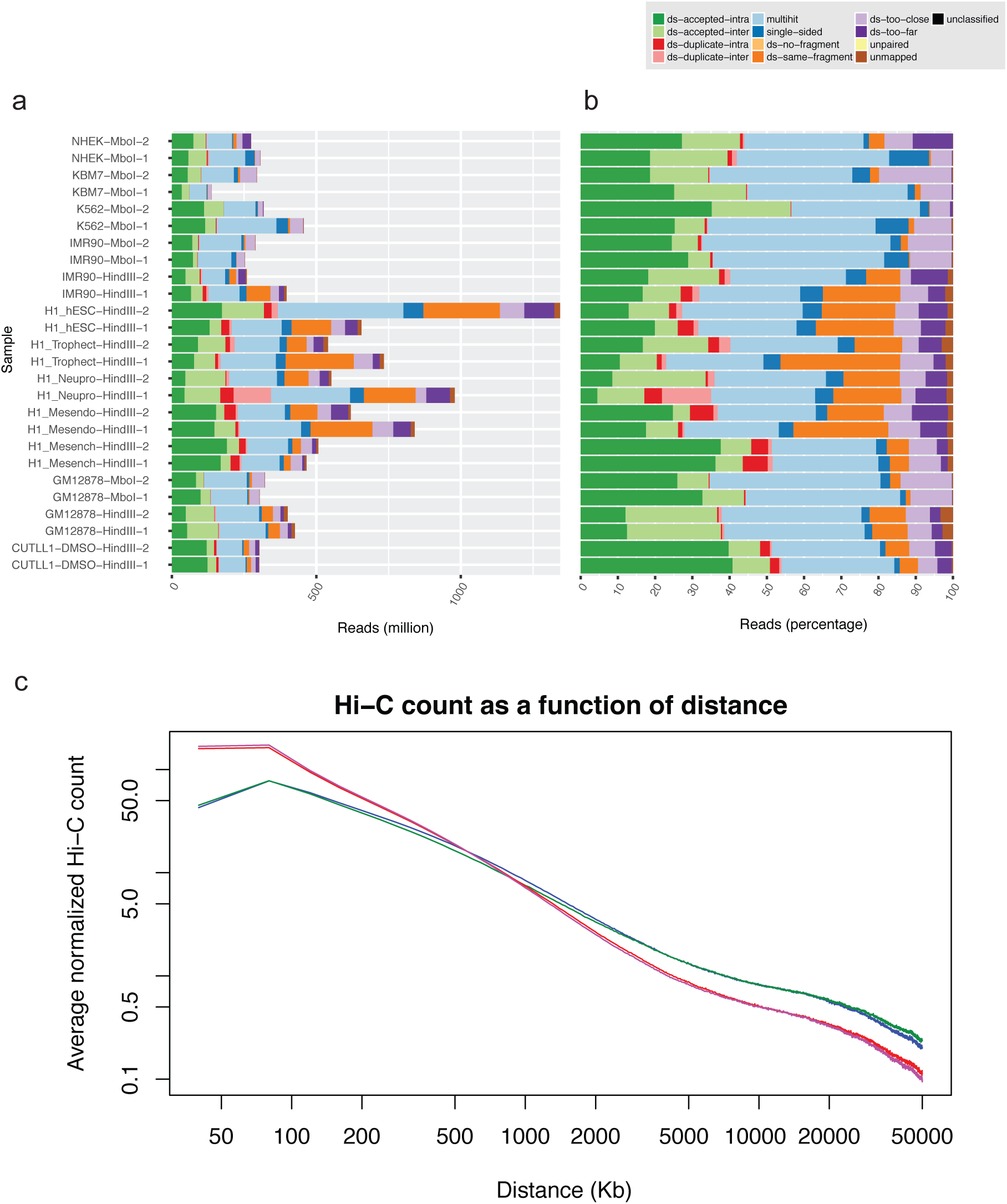
Quality assessment of Hi-C datasets. **(a)** Counts of Hi-C read pairs in various read categories: dark and light green indicate read pairs that were not designated as artifacts and can be used in downstream analyses, **(b)** Percentages of Hi-C reads in each category, **(c)** Average scaled Hi-C read pair count as a function of distance between interacting loci.

**Supplementary Figure 2.**
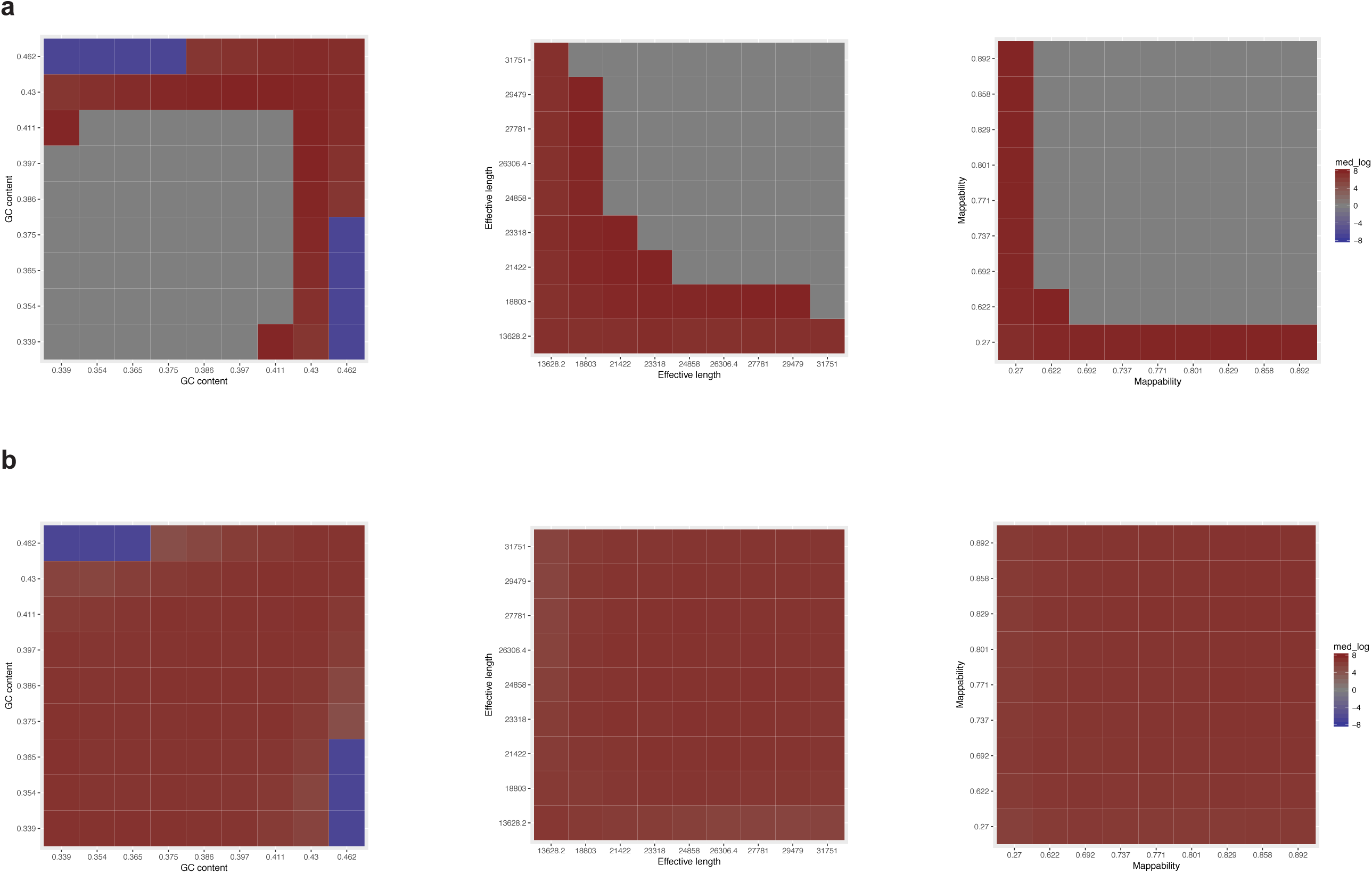
Scaling method corrects for enzyme-specific biases. **(a)** Unprocessed Hi-C matrices exhibit enzyme-specific biases related to the GC content, effective length and mappability of the interacting bins, **(b)** In contrast, our scaling method corrects for all three enzyme-specific biases.

**Supplementary Figure 3.**
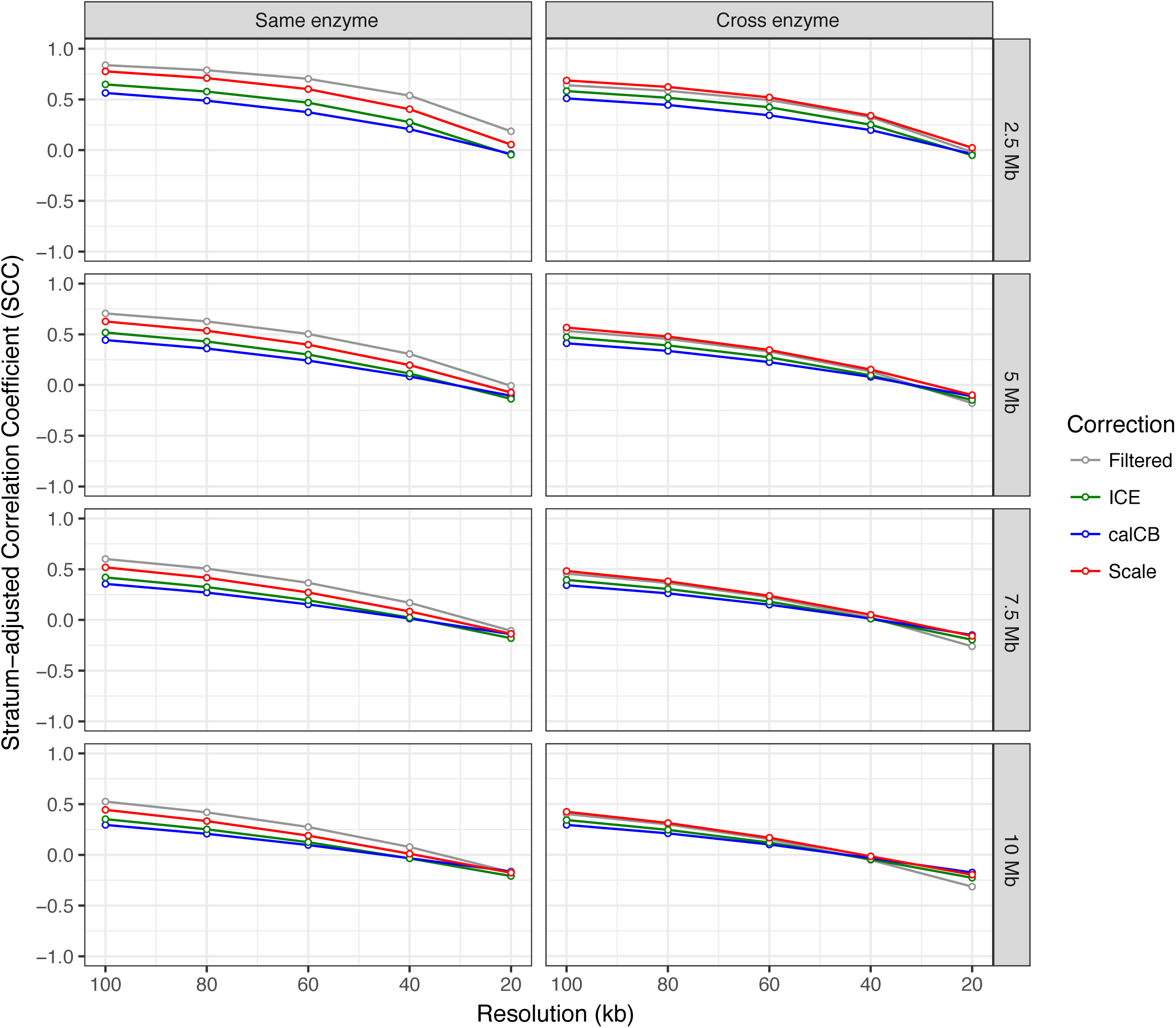
Reproducibility of Hi-C contact matrices *(low sequencing depth)* Comparison of Hi-C contact matrices between biological replicates generated from Hi-C library using the same or different restriction enzymes; Hi-C matrices were estimated using four methods (naïve filtering, iterative correction, calCB and simple scaling); assessment was performed using stratum-adjusted correlation coefficient at resolutions ranging from 100kb to 20kb and maximum distances of 2.5Mb, 5Mb, 7.5Mb and 10Mb between interacting pairs.

**Supplementary Figure 4.**
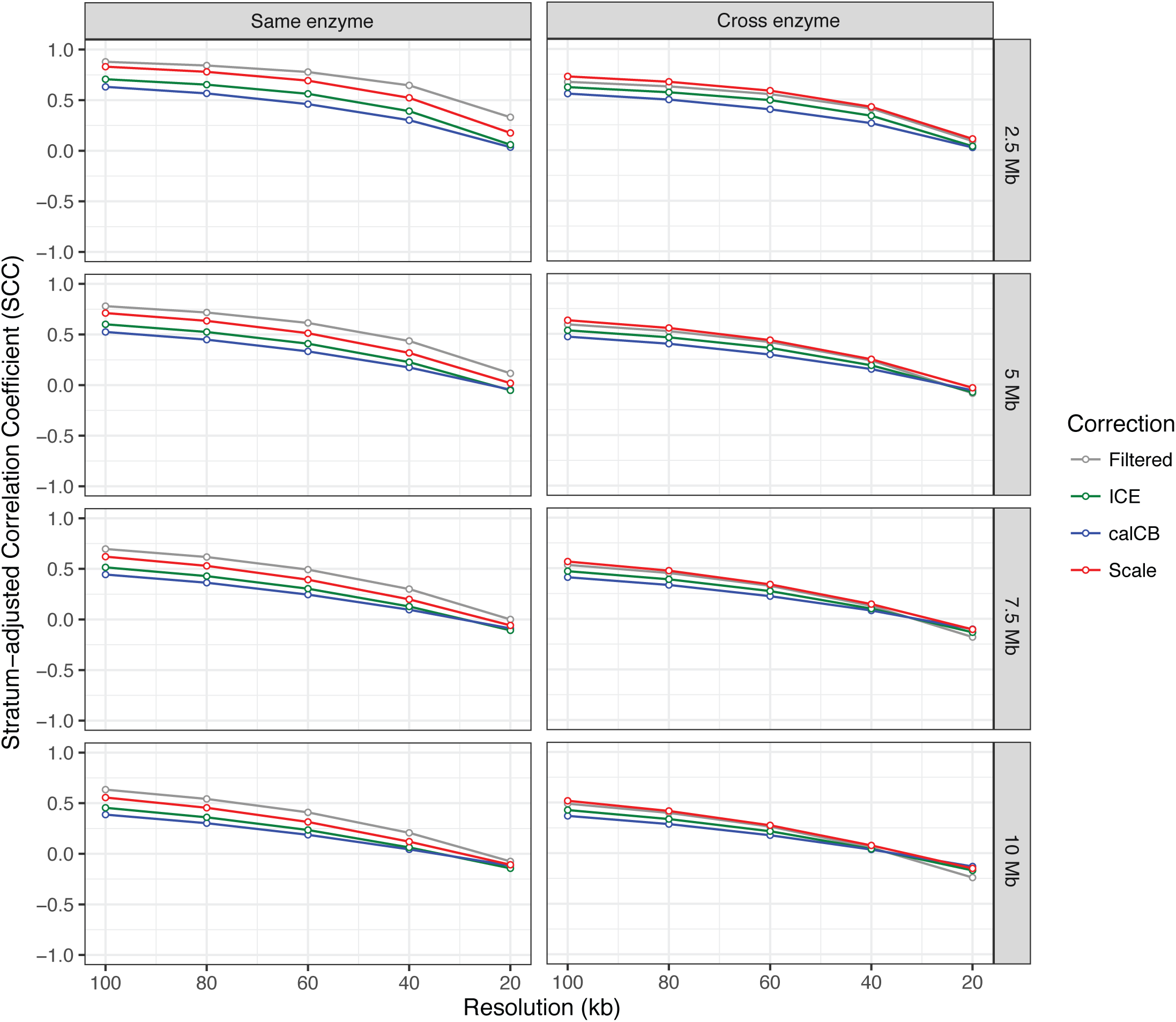
Reproducibility of Hi-C contact matrices *(high sequencing depth)* Comparison of Hi-C contact matrices between biological replicates generated from Hi-C library using the same or different restriction enzymes; Hi-C matrices were estimated using four methods (naïve filtering, iterative correction, calCB and simple scaling); assessment was performed using stratum-adjusted correlation coefficient at resolutions ranging from 100kb to 20kb and maximum distances of 2.5Mb, 5Mb, 7.5Mb and 10Mb between interacting pairs.

**Supplementary Figure 5.**
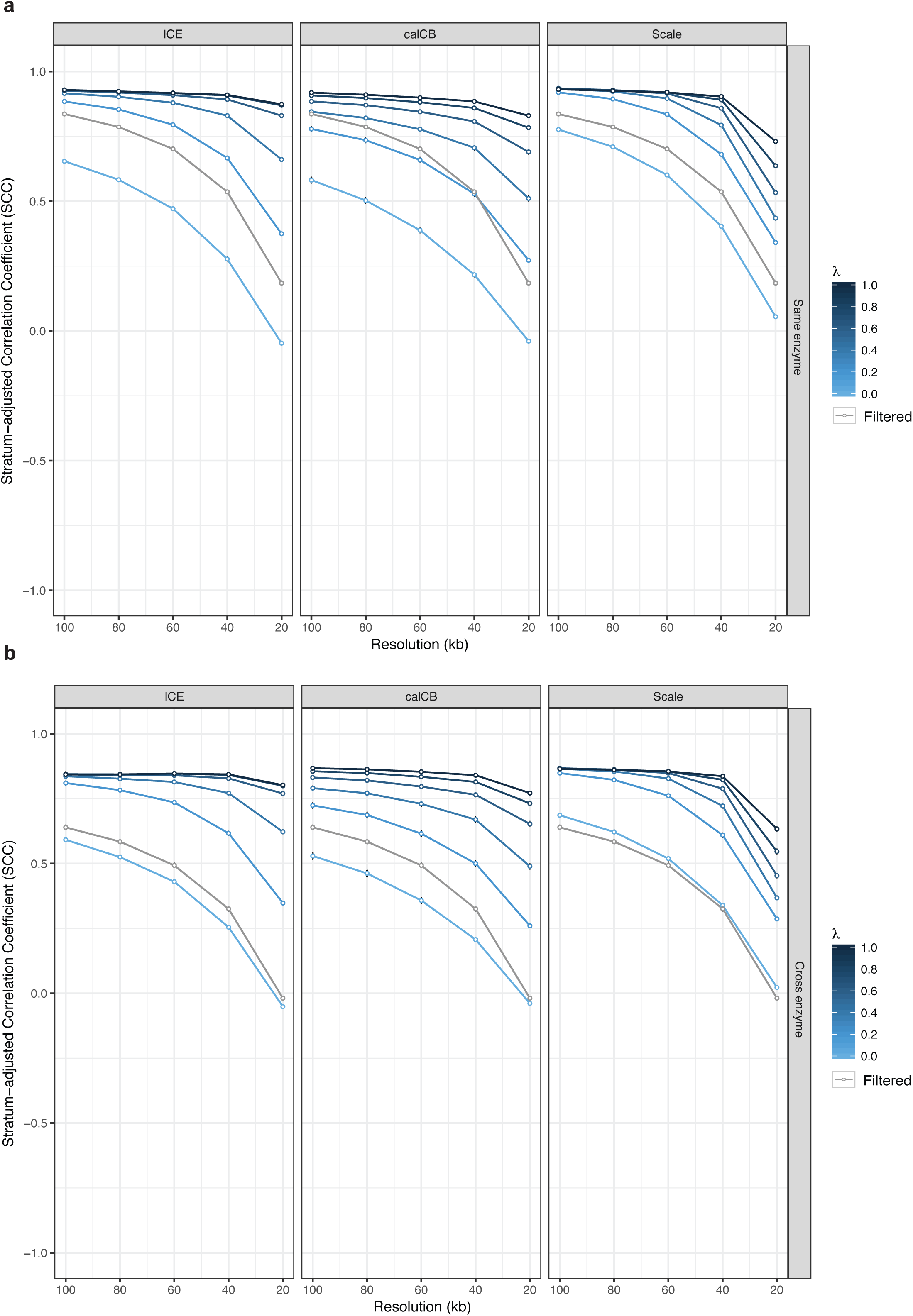
Fused two-dimensional lasso improves reproducibility of Hi-C contact matrices *(low sequencing depth).* **(a)** Stratum-adjusted correlation coefficient values are improved by increasing the value of fused lasso parameter *λ* for matrices estimated with ICE, calCB and our simple scaling method (same enzyme), **(b)** Stratum-adjusted correlation coefficient values are improved by increasing the value of fused lasso parameter *λ* for matrices estimated with ICE, calCB and our simple scaling method (cross enzyme). As a baseline control, stratum-adjusted correlation coefficients of Hi-C contact matrices generated by the naïve filtering method are marked by the gray line in each panel.

**Supplementary Figure 6.**
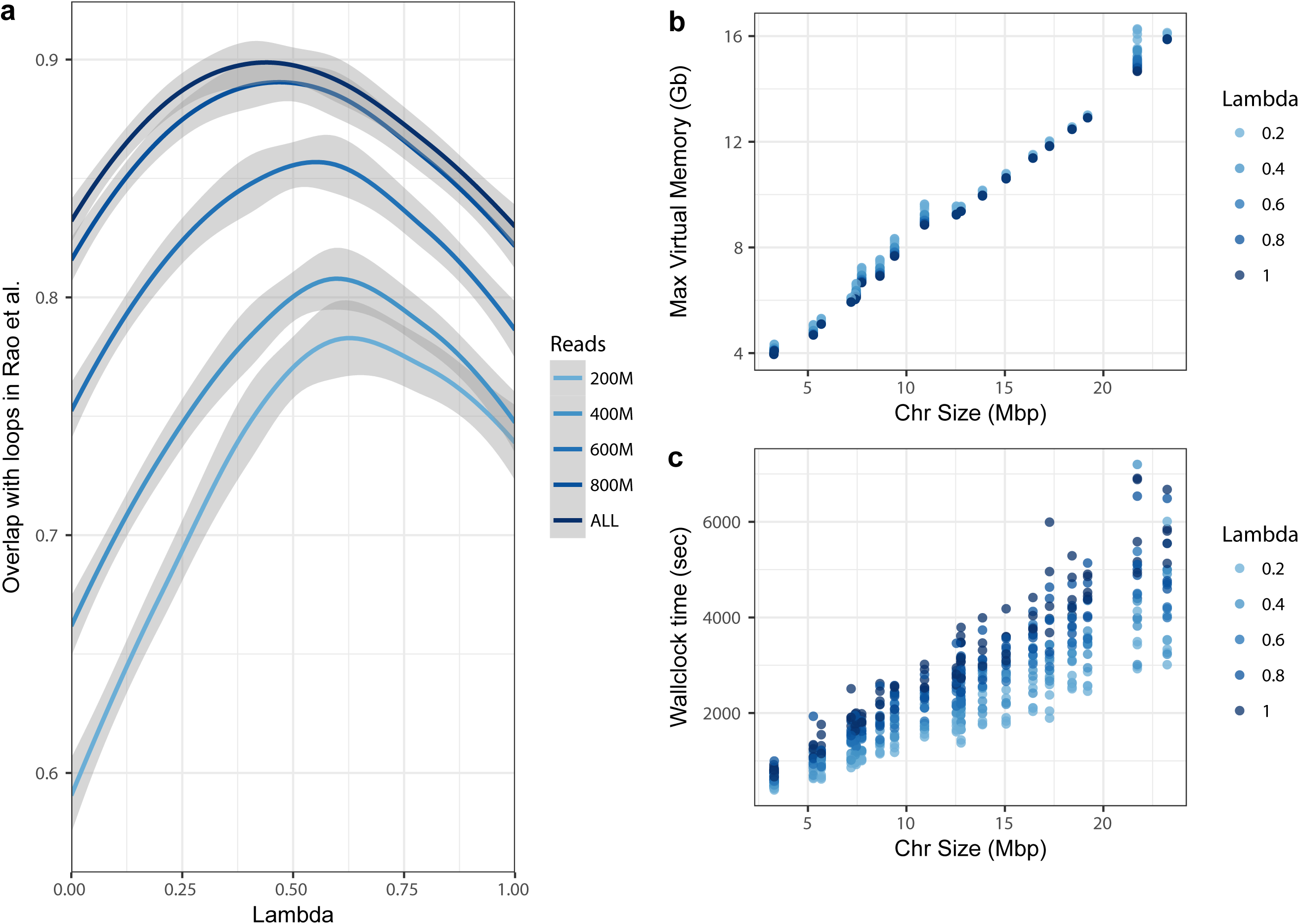
Application of fused 2D lasso at fine resolutions. **(a)** Fraction of loops reported in [20] at 5kb resolution that are within the top 10% of DNA-DNA interactions obtained from the application of the fused lasso algorithm (for different *λ* values and varying sequencing depth) on the ICE-corrected Hi-C matrices obtained from [20], **(b)** Maximum memory requirements of the fused lasso algorithm as a function of chromosome size, **(c)** CPU usage of the fused lasso algorithm as a function of chromosome size.

**Supplementary Figure 7.**
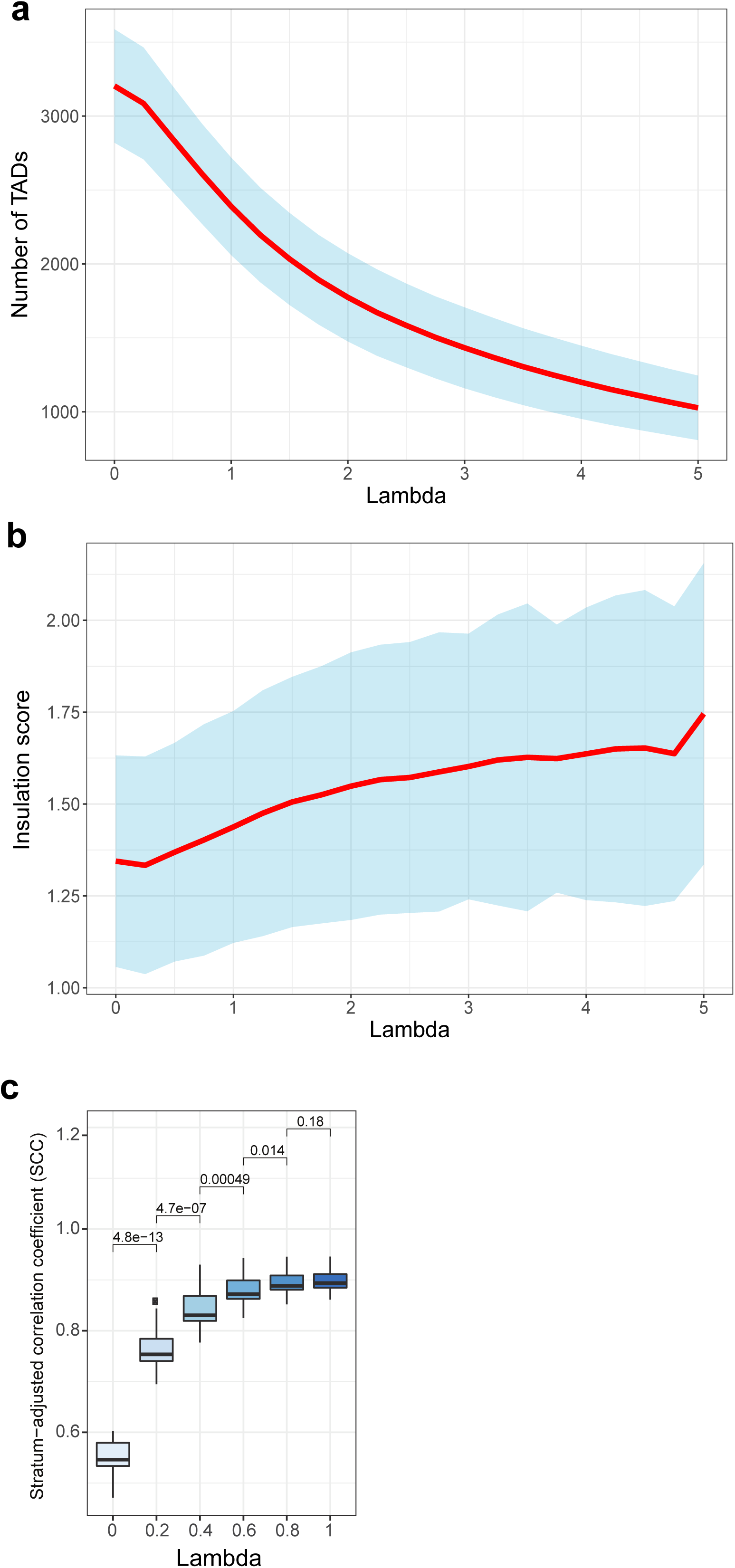
Association of λ-boundaries with insulation score. **(a)** Number of TADs for *λ* values ranging from 0 to 5, **(b)** TAD boundaries lost at higher values of parameter *λ* are associated with higher TAD insulation scores, **(c)** Calculation of optimal *λ* by comparing the statistical significance (measured by a two-sided Wilcoxon rank sum test) of the improvement of reproducibility between successive *λ* values (each dot in the boxplots represents a different chromosome).

**Supplementary Figure 8.**
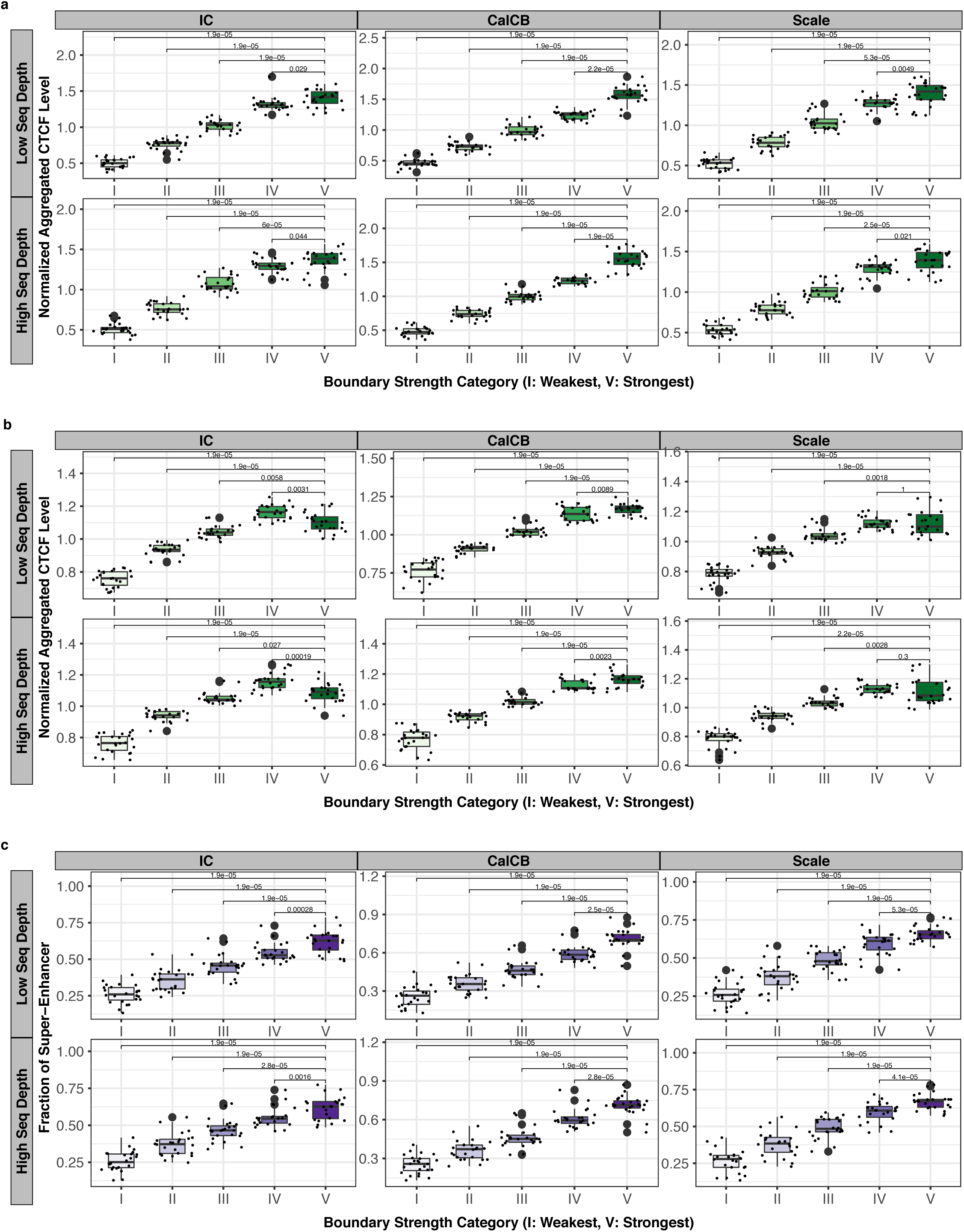
Robustness analysis of the classification and characterization of TAD boundaries according to the “ratio” insulation score. Independent of Hi-C matrix “correction” method and sequencing depth, TAD boundary “ratio” insulation scores are associated with: **(a)** CTCF levels at TSSs, **(b)** CTCF levels (both TSS and non-TSS), **(c)** proximity to super-enhancer elements.

**Supplementary Figure 9.**
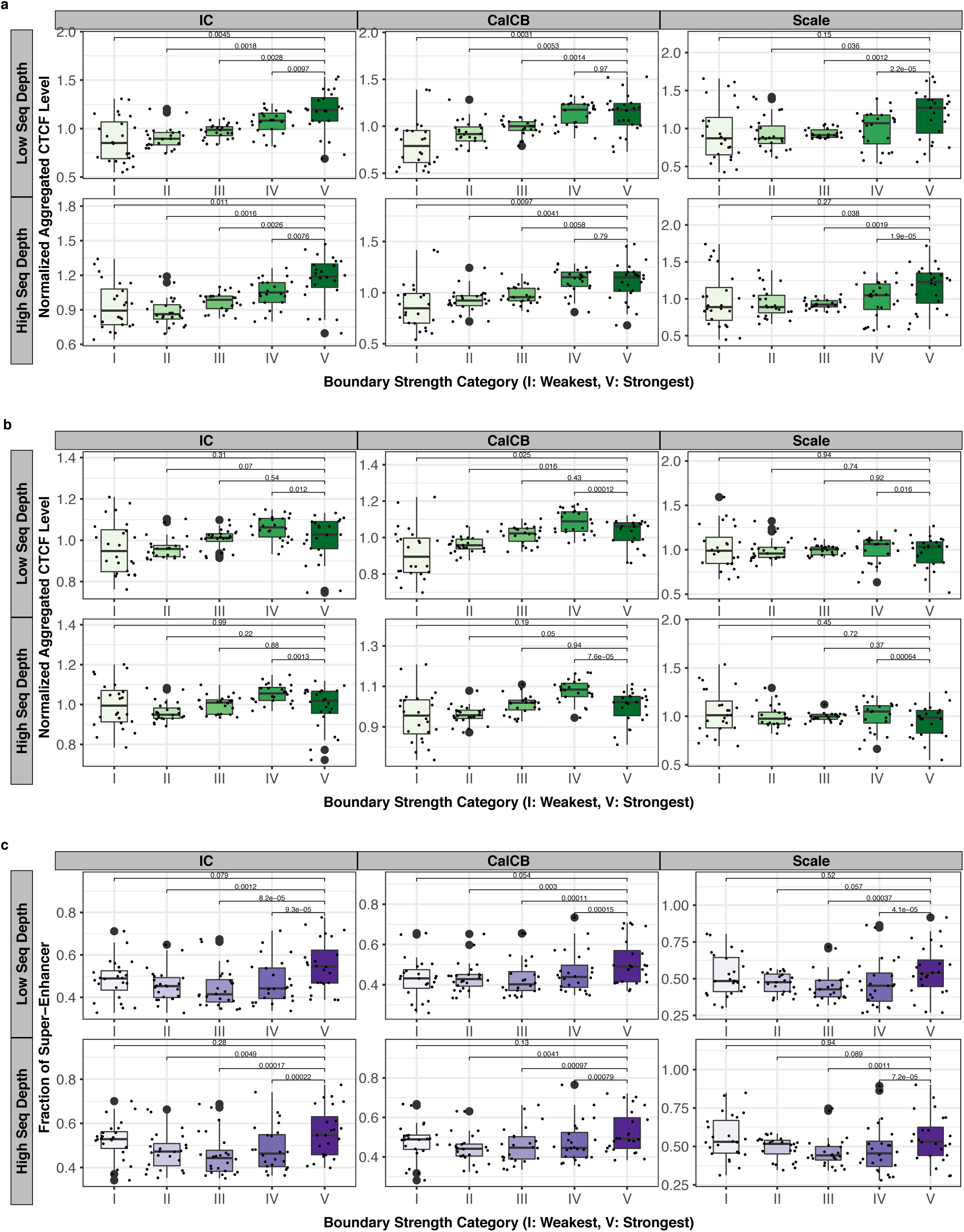
Robustness analysis of the classification and characterization of TAD boundaries according to the “crane” insulation score. Independent of Hi-C matrix “correction” method and sequencing depth (but not as significantly as “ratio” insulation scores) TAD boundary “crane” insulation scores are (weakly) associated with: **(a)** CTCF levels at TSSs, **(b)** CTCF levels (both TSS and non-TSS), **(c)** proximity to super-enhancer elements.

**Supplementary Figure 10.**
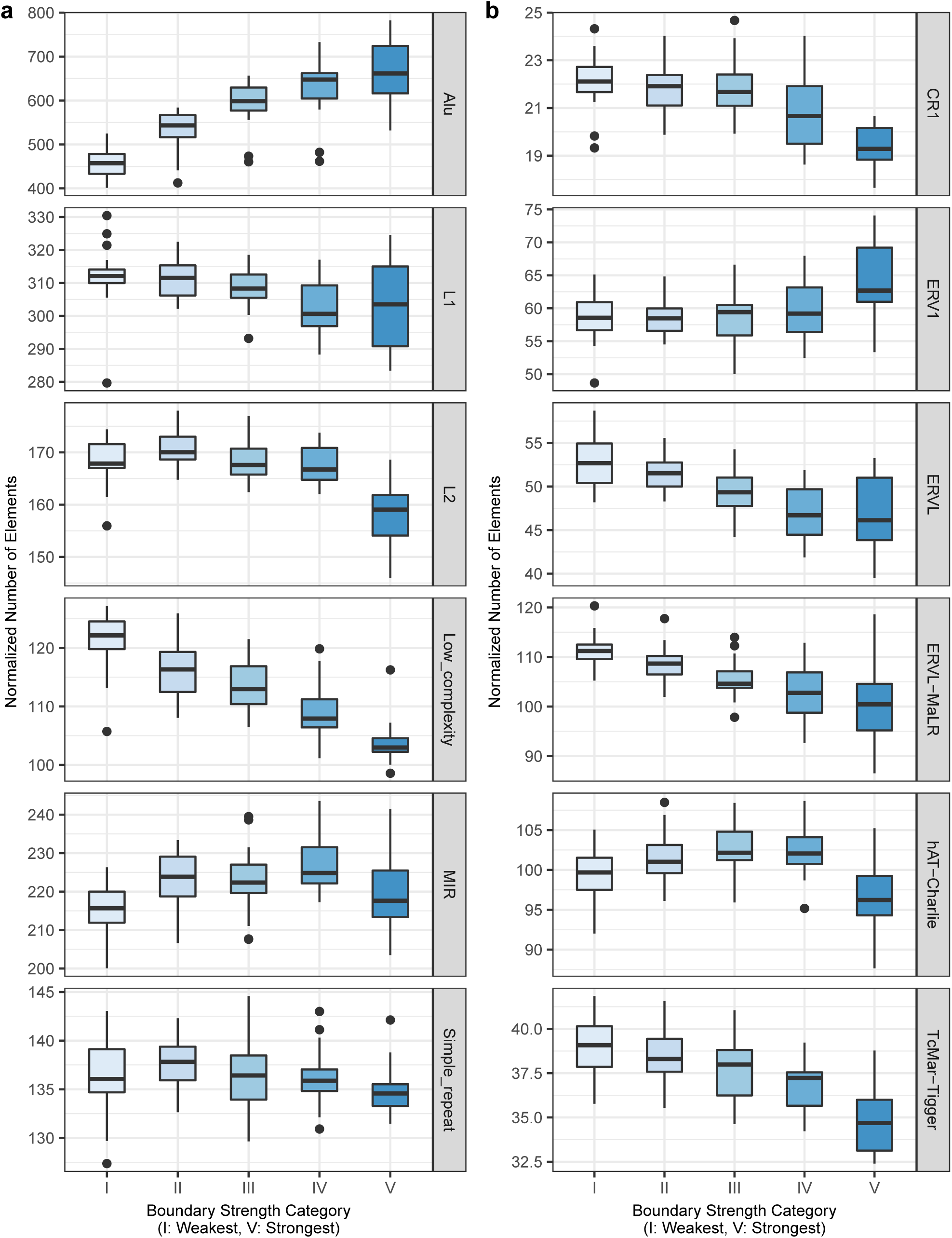
Association of repeat elements with TAD insulation scores. Normalized numbers of repeat elements in proximity to boundaries of certain boundary strength. The distributions represented by the boxplots are across samples and replicates. Darker blue in the blue colour gradient denotes higher boundary strength.

**Supplementary Figure 11.**
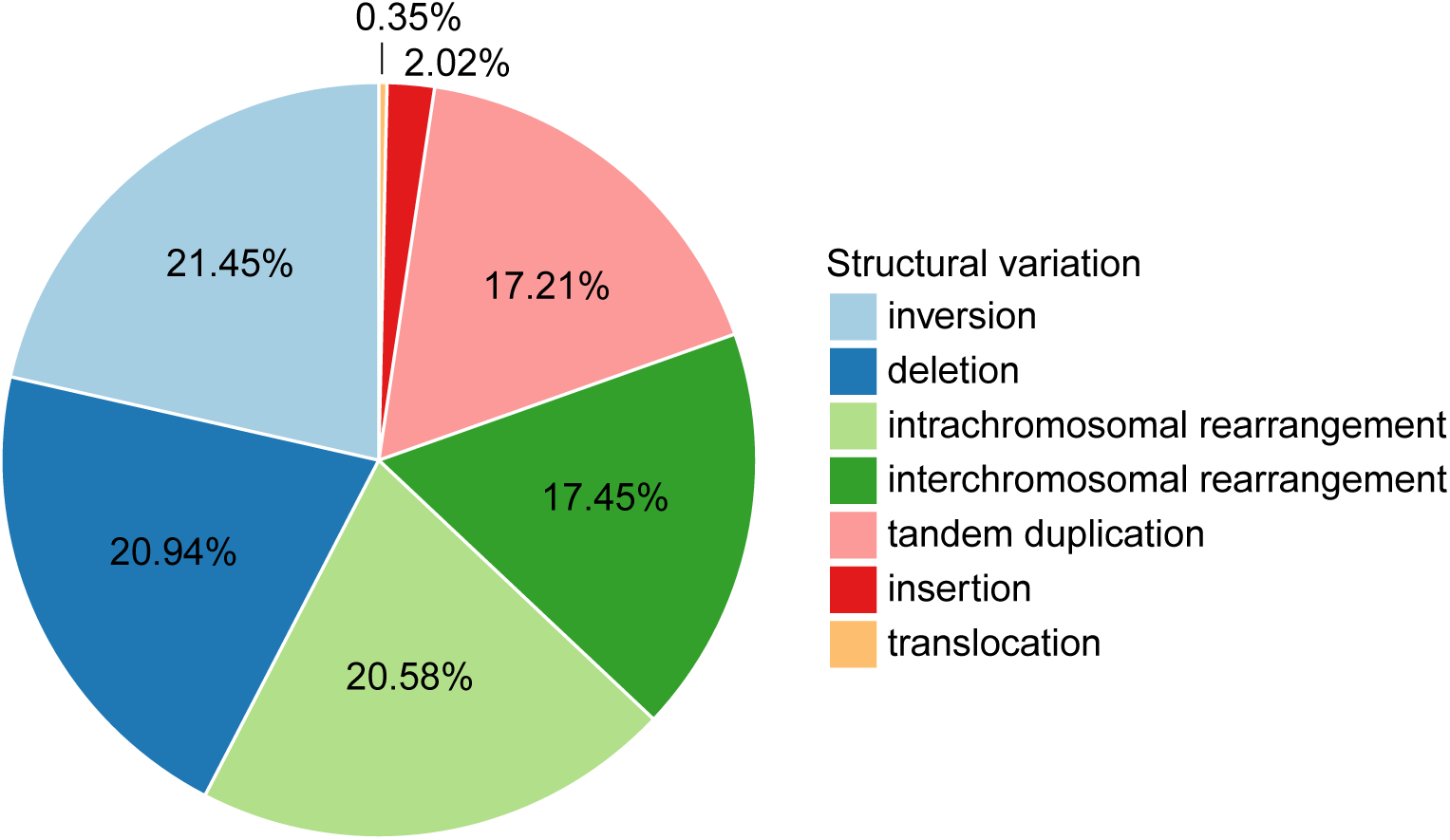
Distribution of somatic structural alterations in the ICGC database.

**Supplementary Figure 12.**
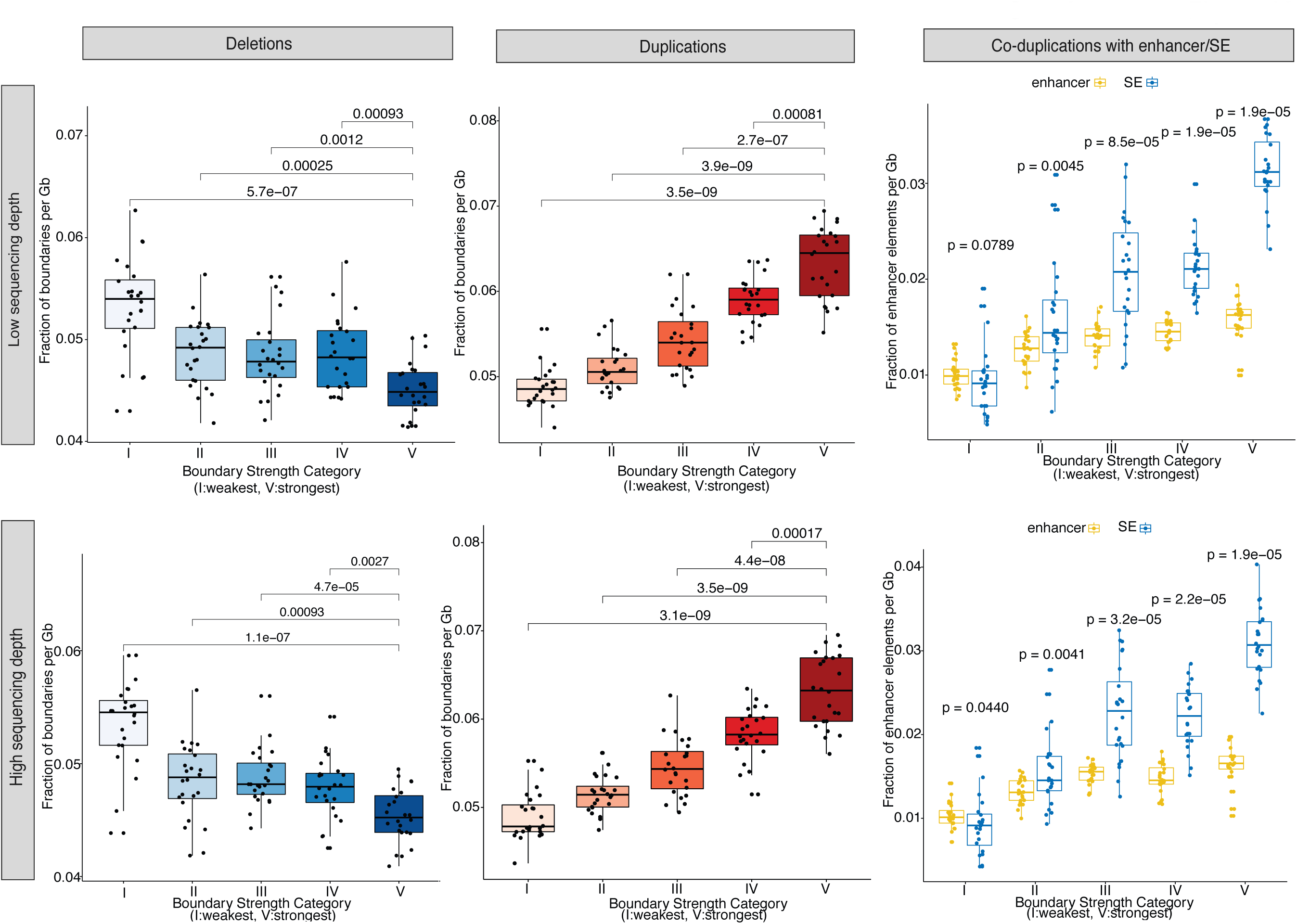
Robustness analysis of the association of strong vs weak TAD boundaries with deletions and tandem duplications in cancer patients. Results presented in **Figure 5a-c** are consistent, independent of sequencing depth: **(a)** Classification of focally deleted boundaries in cancer according to their strength, **(b)** Classification of focally duplicated boundaries in cancer according to their strength, **(c)** Co-duplications of TAD boundaries with regular enhancers and super-enhancers in cancer.

**Supplementary Table 1. Description and accession numbers of Hi-C, CTCF and H3K27ac datasets.**

